# Comparative analysis of cell-cell communication at single-cell resolution

**DOI:** 10.1101/2022.02.04.479209

**Authors:** Aaron J. Wilk, Alex K. Shalek, Susan Holmes, Catherine A. Blish

**Affiliations:** Stanford Immunology Program, Stanford University School of Medicine, Stanford, CA 94305, USA; Department of Medicine, Stanford University School of Medicine, Stanford, CA 94305, USA; Medical Scientist Training Program, Stanford University School of Medicine, Stanford, CA 94305, USA; Institute for Medical Engineering & Science, Massachusetts Institute of Technology, Cambridge, MA 02139, USA; Department of Chemistry, Massachusetts Institute of Technology, Cambridge, MA 02139, USA; Koch Institute for Integrative Cancer Research, Massachusetts Institute of Technology, Cambridge, MA 02139, USA; Ragon Institute of MGH, MIT, and Harvard, Cambridge, MA 02139, USA; Broad Institute of MIT and Harvard, Cambridge, MA 02142, USA; Department of Statistics, Stanford University, Stanford, CA 94305, USA; Chan Zuckerberg Biohub, San Francisco, CA 94158, USA

**Author notes:** These authors jointly supervised this work (Susan Holmes, Catherine A. Blish).

**Keywords:** cell-cell communication, intercellular signaling, single-cell transcriptomics

## Abstract

Inference of cell-cell communication (CCC) from single-cell RNA-sequencing data is a powerful technique to uncover putative axes of multicellular coordination, yet existing methods perform this analysis at the level of the cell type or cluster, discarding single-cell level information. Here we present Scriabin – a flexible and scalable framework for comparative analysis of CCC at single-cell resolution. We leverage multiple published datasets to show that Scriabin recovers expected CCC edges and use spatial transcriptomic data, genetic perturbation screens, and direct experimental manipulation of receptor-ligand interactions to validate that the recovered edges are biologically meaningful. We then apply Scriabin to uncover co-expressed programs of CCC from atlas-scale datasets, validating known communication pathways required for maintaining the intestinal stem cell niche and revealing species-specific communication pathways. Finally, we utilize single-cell communication networks calculated using Scriabin to follow communication pathways that operate between timepoints in longitudinal datasets, highlighting bystander cells as important initiators of inflammatory reactions in acute SARS-CoV-2 infection. Our approach represents a broadly applicable strategy to leverage single-cell resolution data maximally toward uncovering CCC circuitry and rich niche-phenotype relationships in health and disease.

## INTRODUCTION

Complex multicellular organisms rely on coordination within and between their tissue niches to maintain homeostasis and appropriately respond to internal and external perturbations. This coordination is achieved through cell-cell communication (CCC), whereby cells send and receive biochemical and physical signals that influence cell phenotype and function^1,2^. A fundamental goal of systems biology is to understand the communication pathways that enable tissues to function in a coordinated and flexible manner to maintain health and fight disease^3,4^.

The advent of single-cell RNA-sequencing (scRNA-seq) has made it possible to dissect complex multicellular niches by applying the comprehensive nature of genomics at the “atomic” resolution of the single cell. Concurrently, the assembly of protein-protein interaction databases^5^ and the rise of methods for pooled genetic perturbation screening^6,7^ have empowered the development of methods that infer putative axes of cell-to-cell communication from scRNA-seq datasets^8–13^. These techniques generally function by aggregating ligand and receptor expression values for groups of cells to infer which groups of cells are likely to interact with one another^14–17^. However, biologically, CCC does not operate at the level of the group; rather, such interactions take place between individual cells. There exists a need for methods of CCC inference that: 1. analyze interactions at the level of the single cell; 2. leverage the full information content contained within scRNA-seq data by looking at up- and down-stream cellular activity; 3. enable comparative analysis between conditions; and, 4. are robust to multiple experimental designs.

Here we introduce <u>s</u>ingle-<u>c</u>ell resolved interaction <u>a</u>nalysis through <u>bin</u>ning (Scriabin) – an adaptable and computationally-efficient method for CCC analysis. Scriabin dissects complex communicative pathways at single-cell resolution by combining curated ligand-receptor interaction databases^13,18,19^, models of downstream intracellular signaling^20^, anchor-based dataset integration^21^, and gene network analysis^22^ to recover biologically meaningful CCC edges at single-cell resolution.

## RESULTS

### A flexible framework for comparative CCC analysis at single-cell resolution

Our goal is to develop a scalable and statistically robust method for the comprehensive analysis of CCC from scRNA-seq data. Scriabin implements three separate workflows depending on dataset size and analytical goals (**Figure 1**): 1. the cell-cell interaction matrix workflow, optimal for smaller datasets, analyzes communication for each cell-cell pair in the dataset; 2. the summarized interaction graph workflow, designed for large comparative analyses, identifies cell-cell pairs with different total communicative potential between samples; and, 3) the interaction program discovery workflow, suitable for any dataset size, finds modules of co-expressed ligand-receptor pairs.

**Figure 1:**
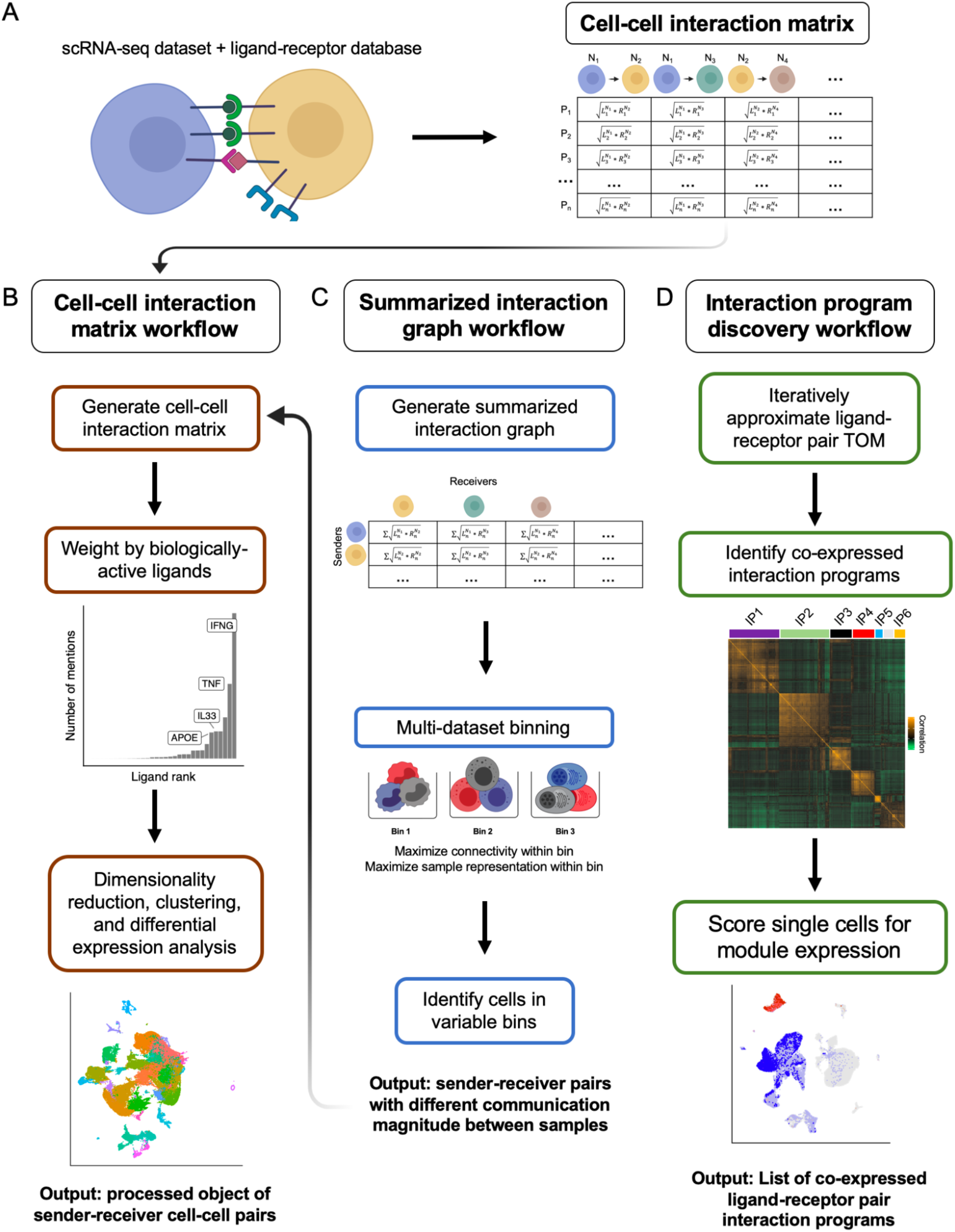
Schematic overview of cell-resolved communication analysis with Scriabin. Scriabin consists of multiple analysis workflows depending on dataset size and the user’s analysis goals. **A)** At the center of these workflows is the calculation of the cell-cell interaction matrix ***M***, which represents all ligand-receptor expression scores for each pair of cells. **B)** <u>Cell-cell interaction matrix workflow</u>: In small datasets, ***M*** can be calculated directly, active CCC edges predicted using NicheNet^20^, and the weighted cell-cell interaction matrix used for downstream analysis tasks like dimensionality reduction. ***M*** is a matrix of NxN cells by P ligand-receptor pairs, where each unique cognate ligand-receptor combination constitutes a unique P. **C)** <u>Summarized interaction graph workflow</u>: In large comparative analyses, a summarized interaction graph ***S*** can be calculated in lieu of a full-dataset ***M***. After high-resolution dataset alignment through binning, the most highly variable bins in total communicative potential can be used to construct an intelligently subsetted ***M*. D)** <u>Interaction program discovery workflow</u>: Interaction programs of co-expressed ligand-receptor pairs can be discovered through iterative approximation of the ligand-receptor pair topological overlap matrix (TOM). Single cells can be scored for the expression of each interaction program (IP), followed by differential expression and modularity analyses.

The fundamental unit of CCC is a sender cell N_i_ expressing ligands that are received by their cognate receptors expressed by a receiver cell N_j_. Scriabin encodes this information in a cell-cell interaction matrix ***M*** by calculating the geometric mean of expression of each ligand-receptor pair by each pair of cells in a dataset (**Figure 1A**). Scriabin supports the use of 15 different protein-protein interaction databases for defining potential ligand-receptor interactions and by default uses the OmniPath database, as this database contains robust annotation of gene category, mechanism, and literature support for each potential interaction^18,19^. As ligand-receptor interactions are directional, Scriabin considers each cell separately as a “sender” (ligand expression) and as a “receiver” (receptor expression), thereby preserving the directed nature of the CCC network. ***M*** can be treated analogously to a gene expression matrix and used for dimensionality reduction, clustering, and differential analyses.

Next, Scriabin identifies biologically meaningful edges, which we define as ligand-receptor pairs that are predicted to affect observed gene expression profiles in the receiving cell (**Figure 1**). This requires defining a gene signature for each cell that reflects its relative gene expression patterns and determining which ligands are most likely to drive that observed signature. First, variable genes are identified in order to immediately focus the analysis on features that distinguish samples of relevance or salient dynamics. When analyzing a single dataset, this set of genes could be the most highly-variable genes (HVGs) in the dataset, which would likely reflect cell type- or state-specific modes of gene expression. Alternatively, when analyzing multiple datasets, the genes that are most variable between conditions (or time points) could be used. To define the relationship between the selected variable genes and each cell, the single cells and chosen variable genes are placed into a shared low dimensional space with multiple correspondence analysis (MCA) implemented by Cell-ID^23^, a weighted generalization of principal component analysis (PCA) that applies to count data. A cell’s gene signature is defined as the set of genes in closest proximity to the variable genes in the MCA embedding (see **Methods**; **Supplementary Text**). An implementation of NicheNet^20^ is then used to nominate the ligands that are most likely to result in each cell’s observed gene signature. Ligand-receptor pairs that are recovered from this process are used to weight the cell-cell interaction matrix ***M*** proportionally to their predicted activity, highlighting the most biologically important interactions (**Figure 1**).

Because one dimension of ***M*** is N x N cells long, it is impractical to construct ***M*** for samples with high cell numbers; this problem will likely be exacerbated as scRNA-seq platforms continue to increase in throughput. Conceptually, solutions to this problem include subsampling and aggregation. Subsampling, however, is statistically inadmissible because it involves omission of available valid data and introduction of sampling noise^24^; meanwhile, aggregation at any level raises the possibility of obscuring important heterogeneity and/or specificity.

A third solution is to first intelligently identify cell-cell pairs of interest and build ***M*** using only those sender and receiver cells. We hypothesize that, in the context of a comparative analysis, sender-receiver cell pairs that change substantially in their *magnitude* of interaction are the most biologically informative. To identify these cells, Scriabin first constructs a summarized interaction graph ***S***, characterized by an N by N matrix containing the sum of all cognate ligand-receptor pair expression scores for each pair of cells. ***S*** is much more computationally efficient to generate, store, and analyze than a full-dataset ***M*** (for a 1,000 cell dataset, ***S*** is 1,000 by 1,000, whereas ***M*** is ∼3,000 by 1,000,000). Comparing summarized interaction graphs from multiple samples requires that cells from different samples share a set of labels or annotations of cells representing the same identity. We use recent progress in dataset integration methodology^21,25^ to develop a high-resolution alignment process we call “binning,” where we assign each cell a bin identity that maximizes the similarity of cells within each bin, maximizes the representation of all samples we wish to compare within each bin, while simultaneously minimizing the degree of agglomeration required (**Figure 1**; **Supplementary Text**). Sender and receiver cells belonging to the bins with the highest communicative variance can then be used to construct ***M***.

Finally, Scriabin implements a workflow for single cell-resolved CCC analysis that is scalable to any dataset size, enabling discovery of co-expressed ligand-receptor interaction programs. This workflow is motivated by the observation that transcriptionally similar sender-receiver cell pairs will tend to communicate through similar sets of ligand-receptor pairs. To achieve this, we adapted the well-established weighted gene correlation network analysis (WGCNA) pipeline^22^ – designed to find modules of co-expressed genes – to uncover modules of ligand-receptor pairs co-expressed by the same sets of sender-receiver cell pairs, which we call “interaction programs”. Scriabin calculates sequences of ***M*** subsets that are used to iteratively approximate a topological overlap matrix (TOM) which is then used to discover highly connected interaction programs. Because the dimensionality of the approximated TOM is consistent between datasets, this approach is highly scalable. The connectivity of individual interaction programs is then tested for statistical significance, which can reveal differences in co-expression patterns between samples. Single cells are then scored for the expression of statistically significant interaction programs. Comparative analyses include differential expression analyses on identified interaction programs, as well as comparisons of intramodular connectivity between samples.

To illustrate the importance of performing CCC analyses at single-cell resolution, we examined CCC of T cells in the tumor microenvironment. Due to their low RNA content, it is often difficult to infer the functional states of T cells from their transcriptomes^26^; yet, T cells participate in communicative pathways that are important to clinical and therapeutic outcomes^27^. Additionally, transcriptional evidence suggests that helper T cells may exist on a phenotypic continuum rather than in traditional discrete functional archetypes^28^. In a dataset of squamous cell carcinoma (SCC) and matched controls^29^, we found a high degree of whole transcriptome phenotypic overlap between intratumoral T cells and those present in normal skin (**Figure 2A**). Further, though there were exhausted T cells in this dataset, they did not occupy a discrete cluster but were rather equally distributed across multiple clusters (**Figure 2A**; **Supplementary Figure 1**), precluding cluster-based CCC approaches from detecting communication modalities unique to exhausted T cells without *a priori* knowledge. We tested Scriabin’s utility in exposing the heterogeneity of the T cell communicative phenotype by applying the cell-cell interaction matrix workflow to pairs of T cells and *CD1C*^*+*^ dendritic cells (DCs), the most abundant antigen-presenting cell (APC) in this dataset. This revealed both a clear distinction between communication profiles between tumor and matched normal, as well as distinct populations of cell-cell pairs with exhausted T cells (**Figure 2B**). Compared to their non-exhausted counterparts, exhausted T cells communicated with *CD1C*^*+*^ DCs predominantly with exhaustion-associated markers *CTLA4* and *TIGIT* and lost communication pathways involving pro-inflammatory chemokines like *CCL4* and *CCL5* (**Figure 2C**)^30^. This illustrates the communicative heterogeneity that can be missed by agglomerative techniques.

**Figure 2:**
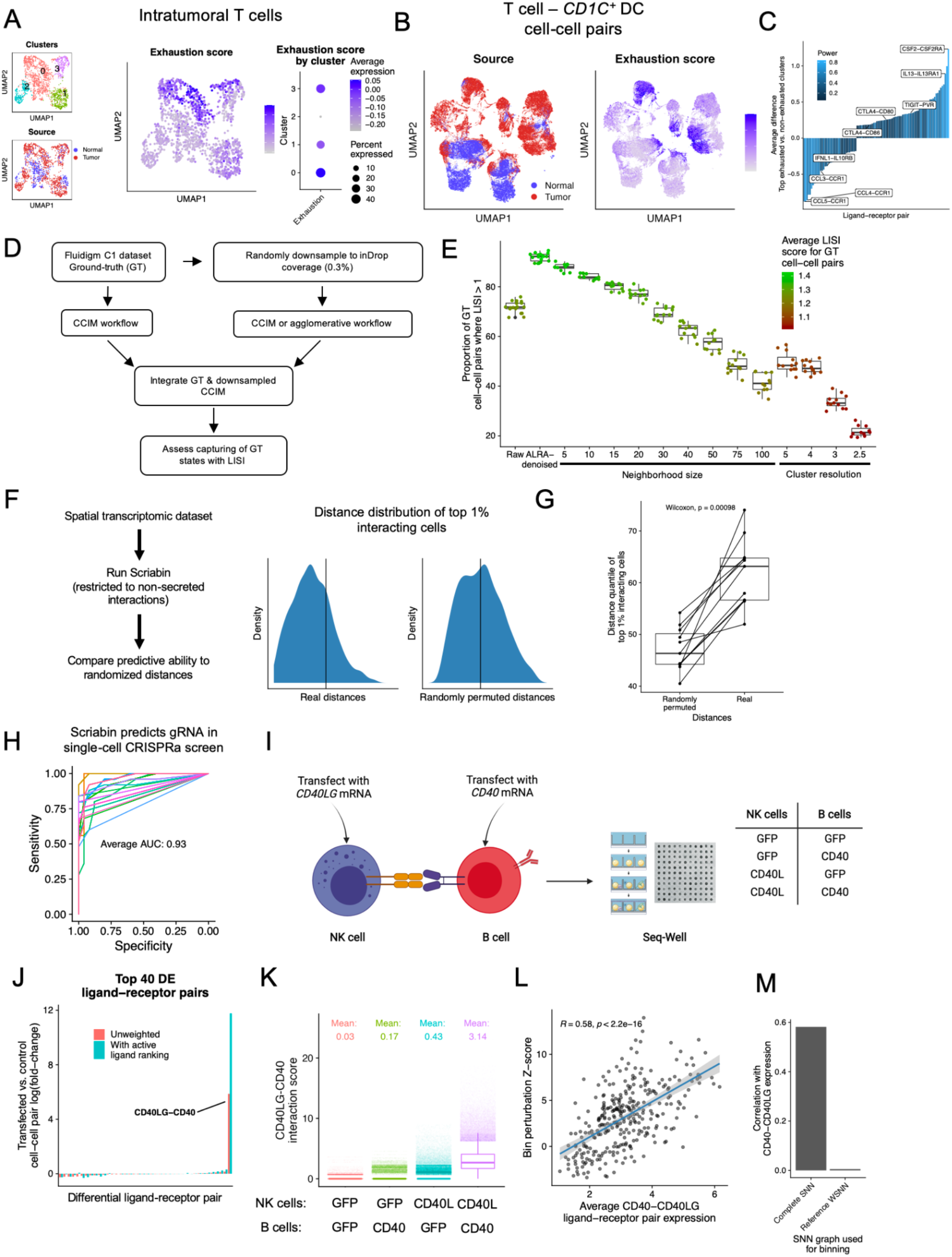
Benchmarking and robustness analysis of cell-resolved communication analysis. **A)** UMAP projections of 1,624 intratumoral T cells from the SCC dataset from Ji, et al.^29^, colored by cluster identity (top left), sample type of origin (either tumor or matched normal; bottom left), and T cell exhaustion score (middle; see **Methods**). The dot plot at right depicts the percent and average expression of the T cell exhaustion score in each cluster. **B)** UMAP projections of 202,708 T cell-*CD1C*^+^ DC cell-cell pairs created from Scriabin’s cell-cell interaction matrix workflow. Points are colored by sample type of origin (left) and the T cell exhaustion score of the T cell in the cell-cell pair (right). **C)** Bar plot depicting differentially-expressed ligand-receptor pairs among T cell-*CD1C*^+^ DC cell-cell pairs between exhausted and non-exhausted T cell senders. Individual bars are colored by the power from Seurat’s implementation of an ROC-based DE test. **D)** Schematic illustrating the workflow to evaluate the impact of technical noise on the robustness of cell-cell communication analyses with Scriabin. **E)** Boxplot depicting the ability of downsampled CCIMs to recapitulate the ground-truth (GT) CCIM. The *y*-axis depicts the proportion of GT cell-cell pairs that are recapitulated by a query cell-cell pair (LISI score > 1), and points are colored by the mean LISI score for GT cell-cell pairs. Each experimental condition listed on the *x*-axis was repeated on 12 different random subsamples of the Fluidigm C1 pancreas islet dataset^21,33^. **F)** Left, description of workflow to validate Scriabin using spatial transcriptomics datasets; right, density plots showing the distribution of cell-cell distances within the top 1% of highly interacting cell-cell pairs predicted by Scriabin. The vertical black lines denote the median distance of all cell-cell pairs. **G)** The procedure depicted in (**F**) was repeated for 11 datasets, and the median distance quantile of the top 1% interacting cell-cell pairs calculated using real cell distances relative to randomly permuted cell distances. Shown is an exact two-sided p-value from the Wilcoxon rank-sum test. **H)** Receiver operating characteristic (ROC) plots depicting Scriabin’s ability to correctly predict the gRNA with which a single cell was transduced based on its communicative profile. Each of the *n* = 15 lines represents a different gene target by gRNAs in a CRISPRa dataset of stimulated T cells^40^. **I)** Experimental scheme to validate Scriabin through transfection of exogenous CCC edges. Briefly, isolated human NK cells were transfected with CD40L-encoding mRNA and isolated human B cells were transfected with CD40-encoding mRNA, followed by 12 hr co-culture and scRNA-seq. In total, 21,538 cells from NK cell-B cell co-cultures were profiled by scRNA-seq. Specific sample sizes for the 4 transfection conditions are: 4,934 (GFP-GFP), 5,665 (GFP-CD40), 4,908 (CD40L-GFP), and 6,031 (CD40L-CD40). **J)** Cell-cell interaction matrices were generated by Scriabin for each co-culture condition with or without ligand activity ranking. The bar plot depicts the top differentially-expressed ligand-receptor pairs between cell-cell pairs from control (GFP/GFP) vs transfected (CD40L/CD40) samples. **K)** Box plot depicting *CD40LG*-*CD40* cell-cell pair interaction scores in each co-culture condition. The *CD40LG*-*CD40* interaction score is derived from cell-cell interaction matrices generated with ligand activity ranking. **L)** Scriabin’s summarized interaction graph workflow was used to bin cells across all four co-culture conditions and to identify the bins with the highest communicative perturbation. The scatter plot depicts the relationship between the *CD40LG*-*CD40* interaction score and the CCC perturbation Dunn Z-test statistic for each of 311 bin-bin pairs (see **Methods**). Pearson’s correlation coefficient, exact p-value, and a 95% confidence interval are shown. **M)** Bar plot depicting the Pearson correlation coefficient between bin perturbation and *CD40LG*-*CD40* interaction score using a full-transcriptome SNN graph for binning compared to a reference-based weighted SNN that does not contain structure related to transfection. For (**G**) and (**K**), boxplot features: minimum whisker, 25th percentile - 1.5 × IQR or the lowest value within; minimum box, 25th percentile; center, median; maximum box, 75th percentile; maximum whisker, 75th percentile + 1.5 × IQR or greatest value within.

### Scriabin is a robust and efficient method for single-cell resolved communication analysis

One potential concern of performing single-cell resolution CCC analysis is that scRNA-seq measurements are inherently sparse and noisy. Aggregative techniques, while frequently obscuring biological heterogeneity, do carry the advantage of utilizing less sparse and therefore more robust expression values. Additionally, using single-cell resolution vs. aggregated pseudobulk measurements for CCC analysis is not a binary option, but rather the ends of an entire spectrum of resolution. Probabilistic denoising techniques for scRNA-seq data^31,32^ use information from transcriptionally similar cells to smooth noise created by putative technical zeroes, and represent a mild form of aggregation by smoothing measured expression values. Further, cluster-based agglomerative CCC techniques can operate at a wide range of potential clustering resolutions. We sought to quantitatively examine the impact of technical noise on single-cell resolution CCC analysis and identify if there is an optimal degree of aggregation that avoids issues with data sparsity without agglomerating over distinct communication phenotypes.

To do this, we simulated technical noise by randomly downsampling a deeply-sequenced scRNA-seq dataset (**Figure 2D**). We defined as ground-truth a dataset of pancreatic islet cells profiled by the Fluidigm C1 platform^21,33^, which profiles cells approximately two orders of magnitude more deeply than droplet-based methods. We then randomly downsampled this dataset to the sequencing depth of inDrop (0.3% of counts relative to Fluidigm C1)^34,35^. We performed Scriabin’s CCIM workflow either directly on this downsampled dataset, on the downsampled dataset denoised by Adaptively thresholded Low-Rank Approximation (ALRA) ^32^, on datasets created by aggregating cells over similarity neighborhoods of nine different sizes, or on pseudobulk expression values from clustering at four different resolutions. Next, we integrated the CCIM generated from the ground-truth Fluidigm C1 dataset with the CCIMs generated from the randomly downsampled dataset. To quantify the degree to which the CCIMs from the downsampled dataset recapitulated the ground-truth CCIM, we calculated the local inverse Simpson’s index (LISI; **Figure 2D**)^36^. This value defines the number of datasets in the neighborhood of each ground-truth cell-cell pair and ranges between 1, denoting that only ground-truth cell-cell pairs are present in the neighborhood, and 2, denoting an equal mixture of ground-truth and downsampled cell-cell pairs.

We found that CCIMs generated from ALRA-denoised data best recapitulated ground-truth data (**Figure 2E**). While downsampling did introduce technical noise when analyzing at single-cell resolution (where ∼70% of ground-truth cell-cell pairs have a downsampled cell-cell pair in their neighborhood), this is almost completely rescued via data denoising. When defining each cell’s transcriptome as the mean transcriptome of that cell and its *k* nearest neighbors, increasing *k* worsened the recapitulation of the ground-truth dataset. ALRA-denoised data outperformed all nine *k* tested. Further, at all cluster resolutions tested, at least 50% of ground-truth CCC states are not captured by using pseudobulk expression values. These data suggest that denoising methods may represent an optimal degree of data aggregation/smoothing that decreases the impacts of technical noise while preserving data structure and heterogeneity. We recommend that users perform denoising when using Scriabin to analyze datasets from platforms with a high degree of sparsity.

We next explored Scriabin’s performance in comparison to other published CCC methods. Scriabin was faster than five agglomerative CCC methods^15–17,37,38^ in analyzing a single dataset at all the dataset sizes tested (**Supplementary Figure 2A**). Of these five agglomerative CCC methods, only Connectome^38^ supports a full comparative workflow, and was slower than Scriabin in a comparative CCC analysis of two datasets (**Supplementary Figure 2B**). We also compared the top CCC edges predicted by these methods^39^ to a pseudobulk version of Scriabin, finding that the top results returned by Scriabin overlapped highly with three of the five published methods analyzed (Connectome, CellChat, and NATMI; **Supplementary Figure 2C**). The remaining two methods (iTALK and SCA) did not have overlapping results with each other or any of the other tested methods (**Supplementary Figure 2C**).

While Scriabin’s results agreed with several published methods, we also sought to demonstrate more directly that these results were biologically correct. We hypothesized that spatial transcriptomic datasets could be leveraged for this purpose, as cells that Scriabin predicts to be highly interacting should be, on average, in closer proximity. We ran Scriabin on 11 spatial transcriptomic datasets, removing secreted ligand-receptor interactions that could operate over a distance from the ligand-receptor database (**Figure 2F**). Cells that Scriabin predicted were the most highly interacting were in significantly closer proximity relative to randomly permuted distances (**Figure 2G**; **Supplementary Figure 3**), indicating that Scriabin can detect spatial features from dissociated data alone.

We next hypothesized that we could leverage a single-cell resolution pooled genetic perturbation screen to validate Scriabin’s ability to identify biologically relevant shifts in cellular communication phenotypes. In an analysis of a CRISPRa Perturb-seq screen of activated human T cells that included guide RNAs (gRNAs) targeting 15 different cell-surface ligands or receptors^40^, we found that Scriabin could accurately predict the gRNA with which a cell was transduced by analyzing cellular CCC profiles (average AUC 0.93; **Figure 2H**).

To provide direct experimental evidence of Scriabin’s ability to detect changes in CCC, we devised an experiment where we transfected isolated natural killer (NK) cells with mRNA encoding CD40L and isolated B cells with mRNA encoding its cognate receptor CD40 (see **Supplementary Text, Figure 2I**). Following co-culture of the transfected cells, we performed scRNA-seq to assess how the forced expression of exogenous CD40 or CD40L impacted CCC. As NK cells do not normally express CD40L, but B cells can express low levels of CD40 at baseline (**Supplementary Figure 4**), we hypothesized that we would observe enhanced communication along the CD40L-CD40 edge only when *CD40LG* was transfected, and that this would be enhanced when both *CD40LG* and *CD40* were transfected. Using Scriabin’s cell-cell interaction matrix workflow, we found that the *CD40LG-CD40* communication edge was the only ligand-receptor pair that was substantially changed in the transfected conditions (**Figure 2J**). This difference was enhanced by incorporating ligand activity weighting into construction of the cell-cell interaction matrix (**Figure 2J**). In line with our predictions, we also found that communication along the *CD40LG-CD40* axis was strongest when NK cells were transfected with *CD40LG* and further increased by transfecting B cells with *CD40* (**Figure 2K**).

Though the aggregative method Connectome^38^ returned *CD40LG-CD40* as a differential communication edge, it also returned 25 other ligand-receptor pairs as statistically significant (**Supplementary Figure 5**). These additional unexpected differential results appeared to be driven by small shifts in expression of very lowly expressed ligands and receptors (**Supplementary Figure 5**). We also used NicheNet alone to identify differentially active ligands between the transfected and untransfected conditions. While CD40L was returned among the top 20 predicted active ligands, NicheNet predicted that FASLG and PTPRC were more differentially-active despite there being little appreciable difference in the expression of these ligands (**Supplementary Figure 5**). This underlines the utility of using information on both relative ligand and receptor expression as well as downstream gene expression changes in performing comparative CCC analyses.

Finally, we used Scriabin’s summarized interaction graph workflow to bin cells from the four transfection conditions and found a significant correlation between the bin perturbation score and the degree to which the cells in each bin were transfected (**Figure 2L**), demonstrating the utility of this workflow in identifying single cells that have the highest degree of communicative perturbation. This correlation was completely abrogated when binning was performed on data structures not related to transfection such as proximity in a reference neighbor graph (**Figure 2M**). These data provide empirical evidence that Scriabin accurately identifies meaningful changes in CCC.

### Scriabin reveals known communicative biology concealed by agglomerative methods

We further evaluated if Scriabin’s single-cell resolution CCC results returned communicating edges that are obscured by agglomerative CCC methods. To this end, we analyzed a publicly-available dataset of a well-characterized tissue niche: the granulomatous response to *Mycobacterium leprae* infection (**Figure 3A**). Granulomas are histologically characterized by infected macrophages and other myeloid cells surrounded by a ring of Th1 T cells^41–43^. These T cells produce IFN-γ that is sensed by myeloid cells; this communication edge between T cells and myeloid cells is widely regarded as the most important interaction in controlling mycobacterial spread^44–46^. Ma, et al. performed scRNA-seq on skin granulomas from patients infected with *Mycobacterium leprae*, the causative agent of leprosy^41^. This dataset includes granulomas from five patients with disseminated lepromatous leprosy (LL) and 4 patients undergoing a reversal reaction (RR) to tuberculoid leprosy, which is characterized by more limited disease and a lower pathogen burden (**Figure 3A**). Analysis of CCC with Scriabin revealed *IFNG* as the most important ligand sensed by myeloid cells in all analyzed granulomas, matching biological expectations (**Figure 3B**). Baseline NicheNet also returned *IFNG* as the most differentially-active ligand in RR granulomas, although with a lesser degree of specificity than Scriabin (**Supplementary Figure 6**).

**Figure 3:**
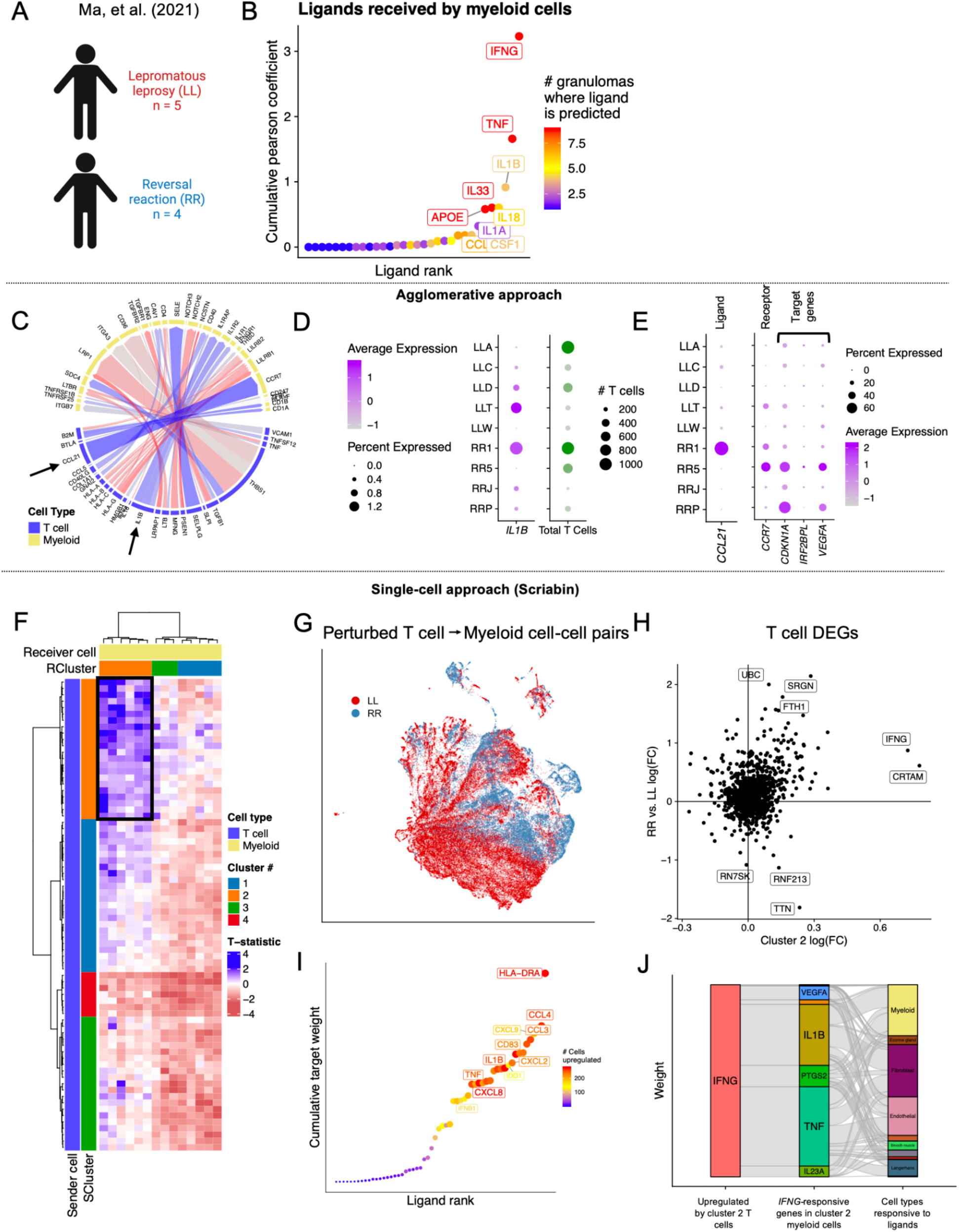
Scriabin reveals communicative pathways obscured by agglomerative techniques. **A)** Schematic of the scRNA-seq dataset of leprosy granulomas published by Ma, et al. Sample sizes for each profiled granuloma are shown in **Supplementary Table 1. B)** Ligands prioritized by Scriabin’s implementation of NicheNet as predicting target gene signatures in granuloma myeloid cells. Points are colored and sized by the number of granulomas in which the ligand is predicted to result in the downstream gene signature. **C)** Circos plot summarizing RR vs. LL differential CCC edges between T cells (senders) and myeloid cells (receivers) generated by Connectome. Blue, edges upregulated in RR; red, edges upregulated in LL. The two black arrows mark T cell-expressed ligands *IL1B* and *CCL21*, which are further analyzed in panels (**D-E**). **D)** Percentage and average of expression of *IL1B* by T cells per granuloma (left), and total number of T cells per granuloma (right). **E)** Percentage and average expression of *CCL21* by T cells per granuloma (left); percentage and average expression of *CCR7* and *CCL21*-stimulated genes by myeloid cells per granuloma. **F)** RR vs. LL differential interaction heatmap between T cell bins (senders; rows) and myeloid cell bins (receivers; columns) generated by Scriabin, colored by the T-statistic between the mean summarized interaction scores of *n* = 4 RR granulomas relative to *n* = 5 LL granulomas. In blue, are bins more highly interacting in RR; in red are the bins more highly interacting in LL. The black box indicates groups of bins predicted to be highly interacting in RR granulomas relative to LL. **G)** UMAP projection of 74,437 perturbed T cell-myeloid cell sender-receiver pairs indicating changes in ligand-receptor pairs used for T cell-myeloid communication in LL vs. RR granulomas. **H)** Scatter plot depicting differential gene expression by T cells. The average log(fold-change) of expression by cluster 2 bins is plotted on the x-axis; the average log(fold-change) of expression by RR granulomas is plotted on the x-axis. **I)** Target genes predicted to be upregulated by *IFNG* in RR granuloma myeloid cells in cluster 2 bins. Points are sized and colored by the number of cells in which the target gene is predicted to be *IFNG*-responsive. **J)** Alluvial plot depicting the RR granuloma cell types that are predicted to receive the *IFNG*-responsive target genes from cluster 2 myeloid cells.

To assess if Scriabin was capable of avoiding pitfalls associated with agglomerative methods in comparative CCC analyses, we analyzed differential CCC pathways from T cells to myeloid cells between LL and RR granulomas using an agglomerative method (Connectome; which implements a full comparative workflow^38^) and Scriabin. We first assessed if it would be possible to analyze higher levels of granularity by using author-provided subclustering annotations. However, as Connectome performs differential CCC analyses by aggregating data at the level of cell type or cluster, this requires that each subcluster have representation from the conditions being compared. In the Ma, et al. dataset, satisfying this condition meant decreasing clustering resolution from 1 to 0.1 so that all subclusters are present in all profiled granulomas and comparing all aggregated LL granulomas to all aggregated RR granulomas (**Supplementary Figure 6**). This requirement moves analysis further from single-cell resolution and we therefore elected to use author-annotated T cells and myeloid cells for analysis without subclustering.

Comparative CCC analysis with Connectome revealed *IL1B* and *CCL21* as the two most upregulated T cell-expressed ligands received by myeloid cells in RR granulomas (**Figure 3C**). However, there was no clear pattern of *IL1B* upregulation among RR granulomas (**Figure 3D**); rather, the RR granuloma that contributed the most T cells expressed the highest level of *IL1B* and the LL granuloma that contributed the most T cells expressed the lowest level of *IL1B* (**Figure 3D**). Additionally, *CCL21* was expressed by T cells of a single RR granuloma, and the myeloid cells of a different RR granuloma expressed the highest levels of the *CCL21* receptor *CCR7* and three *CCL21* target genes (**Figure 3E**). This indicates that the most highly scored differential CCC edges may be due to agglomeration of RR and LL granulomas required by Connectome (**Supplementary Figure 6**), rather than conserved biological changes between these two groups.

To compare differential CCC between LL and RR granulomas with Scriabin, we aligned data from the 9 granulomas together using Scriabin’s binning procedure (**Figure 1**), generated single-cell summarized interaction graphs for each granuloma, and calculated a t-statistic to quantify the difference in interaction for each pair of bins between LL and RR granulomas (**Figure 3F**). This analysis revealed a group of T cell and myeloid bins whose interaction was strongly increased in RR granulomas relative to LL (**Figure 3F**, black box). We visualized the cells in these perturbed bins by generating cell-cell interaction matrices for these cells in each sample and embedding them in shared low dimensional space (**Figure 3G**). The T cells in these bins were defined by expression of *CRTAM*, a marker of cytotoxic CD4 T cells, and upregulated *IFNG* in the RR granulomas (**Figure 3H**). These perturbed T cells were enriched in “RR CTL” and “amCTL” subclusters described by Ma, et al.^41^ that correspond to *IFNG*-expressing cytotoxic T cells (**Supplementary Figure 7**). Perturbed myeloid cells were enriched in transitional macrophage and type I IFN^high^ macrophage subclusters (**Supplementary Figure 7**). Myeloid cells in these bins upregulated several pro-inflammatory cytokines in RR granulomas, including *IL1B, CCL3*, and *TNF* in response to *IFNG* from this T cell subset (**Figure 3I**). *IFNG*-responsive *IL1B* and *TNF* were also predicted to be RR-specific ligands received by myeloid cells, fibroblasts, and endothelial cells in RR granulomas (**Figure 3J**). Collectively, Scriabin identified a subset of *CRTAM*^+^ T cells that upregulated *IFNG* in RR granulomas that is predicted to act on myeloid cells to upregulate additional pro-inflammatory cytokines. These CCC results match previous results demonstrating that enhanced production of *IFNG* can drive RRs^47,48^ and implicate cytotoxic CD4 T cells as initiators of this reaction.

### Discovery of co-expressed interaction programs enables atlas-scale analysis of CCC at single-cell resolution

We next assessed Scriabin’s interaction program discovery workflow. To illustrate the scalability of this process, we chose to analyze a large single-cell atlas of developing fetal gut^49^ composed of 76,592 cells sampled from four anatomical locations (**Figure 4A**). Scriabin discovered a total of 75 significantly correlated interaction programs across all anatomical locations. Scoring all single cells on the expression of the ligands and receptors in these interaction programs revealed strong cell type specific expression patterns for many programs (**Figure 4B**), as well as subtle within-cell type differences in sender or receiver potential, highlighting the importance of maintaining single-cell resolution (**Supplementary Figure 8; Supplementary Text**).

**Figure 4:**
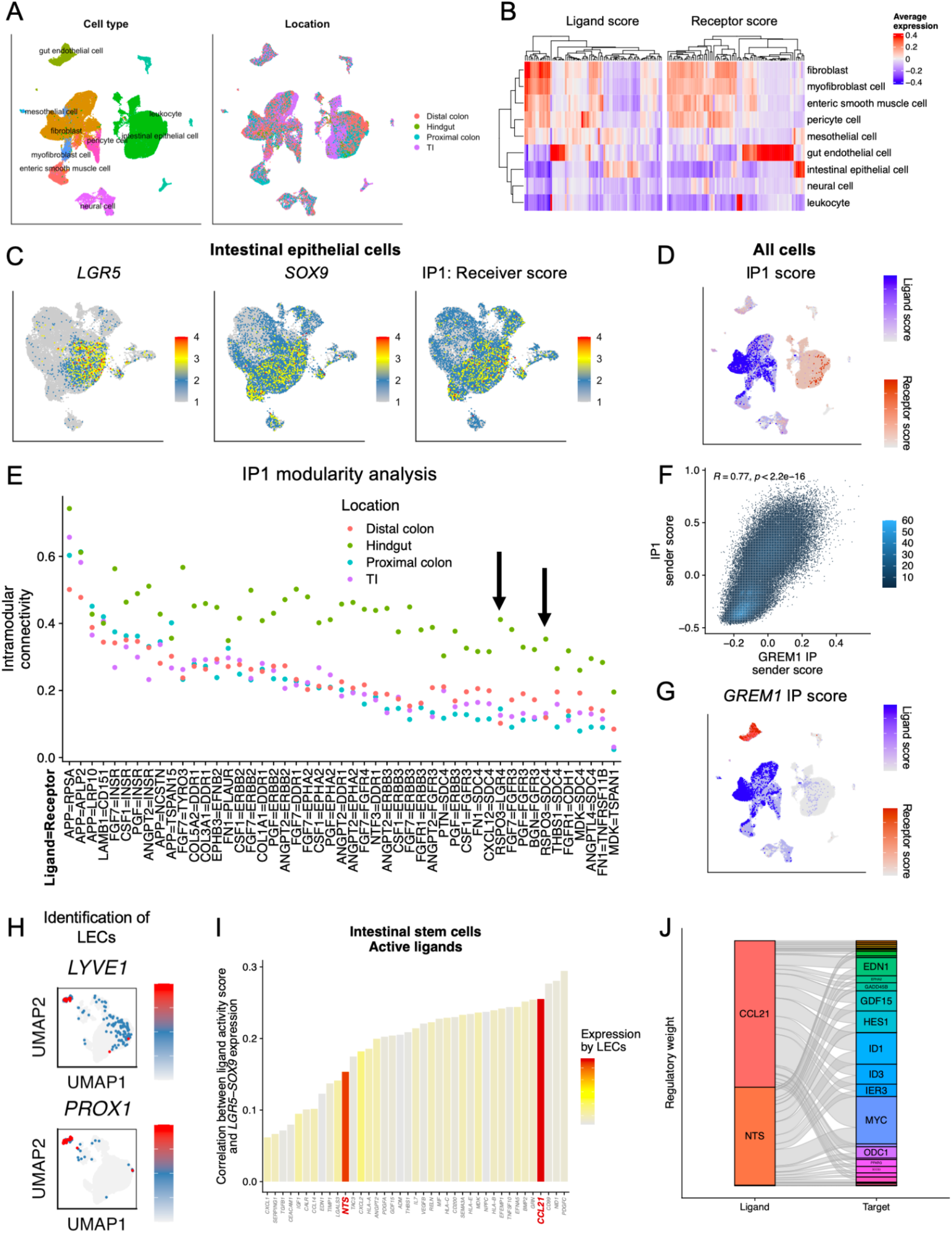
Cell-cell interaction programs of the developing fetal gut. **A**) UMAP projections of the dataset of Fawkner-Corbett, et al.^49^, with 76,592 individual cells colored by author-provided cell type annotations (left), or by anatomical sampling location (right). **B)** Heatmap depicting average expression of interaction program ligands (left) or interaction program receptors (right) by each cell type. **C)** UMAP projections of 25,969 intestinal epithelial cells, colored by expression of stem cell markers *LGR5* and *SOX9*, as well as by the receptor expression score for interaction program 1 (IP1). **D)** UMAP projection of all cells colored by ligand (shades of blue) or receptor (shades of red) expression of IP1. **E)** Intramodular connectivity scores for each ligand-receptor pair in each anatomical location for IP1. The black arrows mark ligand-receptor pairs that include *RSPO3*. **F)** Heatmap of 2d bin counts depicting the correlation between IP1 sender score and the sender score for the IP module that contains the ligand *GREM1*. **G)** UMAP projection of all cells colored by ligand (shades of blue) or receptor (shades of red) expression of the *GREM1* IP. **H)** UMAP projections of 4,447 gut endothelial cells colored by expression of LEC markers *LYVE1* (top) and *PROX1* (bottom). **I)** Bar plot depicting predicted active ligands for intestinal epithelial cells and correlation of predicted ligand activity with expression of ISC markers *LGR5* and *SOX9*. Bars are colored by the average log(fold-change) in expression of each ligand by LECs relative to other gut endothelial cells. **J)** Alluvial plot depicting target genes predicted to be upregulated in ISCs in response to *CCL21* and *NTS*.

We next examined ways in which our identified interaction programs reflected known biology of intestinal development. Recently, several important interactions have been shown to be critical in maintaining the intestinal stem cell (ISC) niche^50–52^. We were able to identify ISCs, defined by expression of *LGR5* and *SOX9*, within the intestinal epithelial cells of this dataset, and discovered a single interaction program (hereafter referred to as IP1) whose receptors were co-expressed with these ISC markers (**Figure 4C**). IP1 represents a program of fibroblast-specific ligand and intestinal epithelial cell receptor expression (**Figure 4D**). Among IP1 ligands were the ephrins *EPHB3*, whose expression gradient is known to control ISC differentiation^53^, and *RSPO3* (**Figure 4E**). Two recent studies have each reported that *RSPO3* production by lymphatic endothelial cells (LECs) and GREM1^+^ fibroblasts is critical for maintaining the ISC niche in mice^51,52^. In this human dataset, we did not observe expression of *RSPO3* in LECs (**Supplementary Figure 8**), and while Fawkner-Corbett, et al. identified *RSPO3* as a potential communication ligand for ISCs^49^, they did not examine the precise source of this ligand. In our application of Scriabin’s interaction program workflow, we found that *GREM1*^+^ fibroblasts expressed *RSPO3* as a part of IP1 that was predicted to be sensed primarily by ISCs, thus demonstrating that this interaction pathway may communicate between different cell types in mouse than in human (**Figure 4D-F**). We also found a separate interaction program containing the ligand *GREM1*; the ligands of this interaction program were co-expressed with IP1 ligands (**Figure 4F**) and predicted to communicate to a different receiver cell type, namely gut endothelial cells (**Figure 4G**).

Despite the absence of *RSPO3* expression in LECs, it remains possible that LECs maintain the ISC niche in human intestinal development, particularly as these cells can reside in close spatial proximity to ISCs^51,52^. While Fawkner-Corbett, et al. included several CCC analyses on endothelial cells, these analyses were performed on aggregated endothelial cells and not specifically on LECs^49^. We were able to identify a small population of LECs (**Figure 4H**), and we examined CCC between LECs and ISCs. We found that two LEC-specific markers, *CCL21* and *NTS*, were predicted to be active ligands for ISCs (**Figure 4I**). *CCL21* and *NTS* were both predicted to result in upregulation of target genes that notably included *MYC* and *ID1* (**Figure 4J**), which are known to participate in intestinal crypt formation and ISC maintenance^54,55^. None of these ligand-receptor CCC edges were returned by an agglomerative CCC analysis by Connectome (**Supplementary Figure 9**). Our results suggest that, unlike in mice, in humans, LECs may contribute to ISC maintenance through production of *CCL21* and *NTS*. Taken together, our results demonstrate the utility of interaction programs both in identifying known CCC edges and providing new biological insights.

### Assembly of longitudinal communicative circuits

A frequent analytical question in longitudinal analyses concerns how events at one time point influence cellular phenotype in the following time point^56,57^. We hypothesized, in datasets with close spacing between time points, that Scriabin’s high-resolution bin identities would allow us to assemble “longitudinal communicative circuits”: chains of sender-receiver pairs across consecutive timepoints. A communicative circuit consists of at least four cells across at least two time points: sender cell at time point 1 (S_1_), receiver cell at time point 1 (R_1_), sender cell at time point 2 (S_2_), and receiver cell at time point 2 (R_2_). If the interaction between S_1_-R_1_ is predicted to result in the upregulation of ligand L_A_ by R_1_, S_1_-R_1_-S_2_-R_2_ participates in a longitudinal circuit if R_1_ and S_2_ share the same bin (i.e., S_2_ represents the counterpart of R_1_ at timepoint 2) and if L_A_ is predicted to be an active ligand in the S_2_-R_2_ interaction (**Figure 5A**). This process enables the stitching together of multiple sequential timepoints to identify communicative edges that are downstream in time and mechanism.

**Figure 5:**
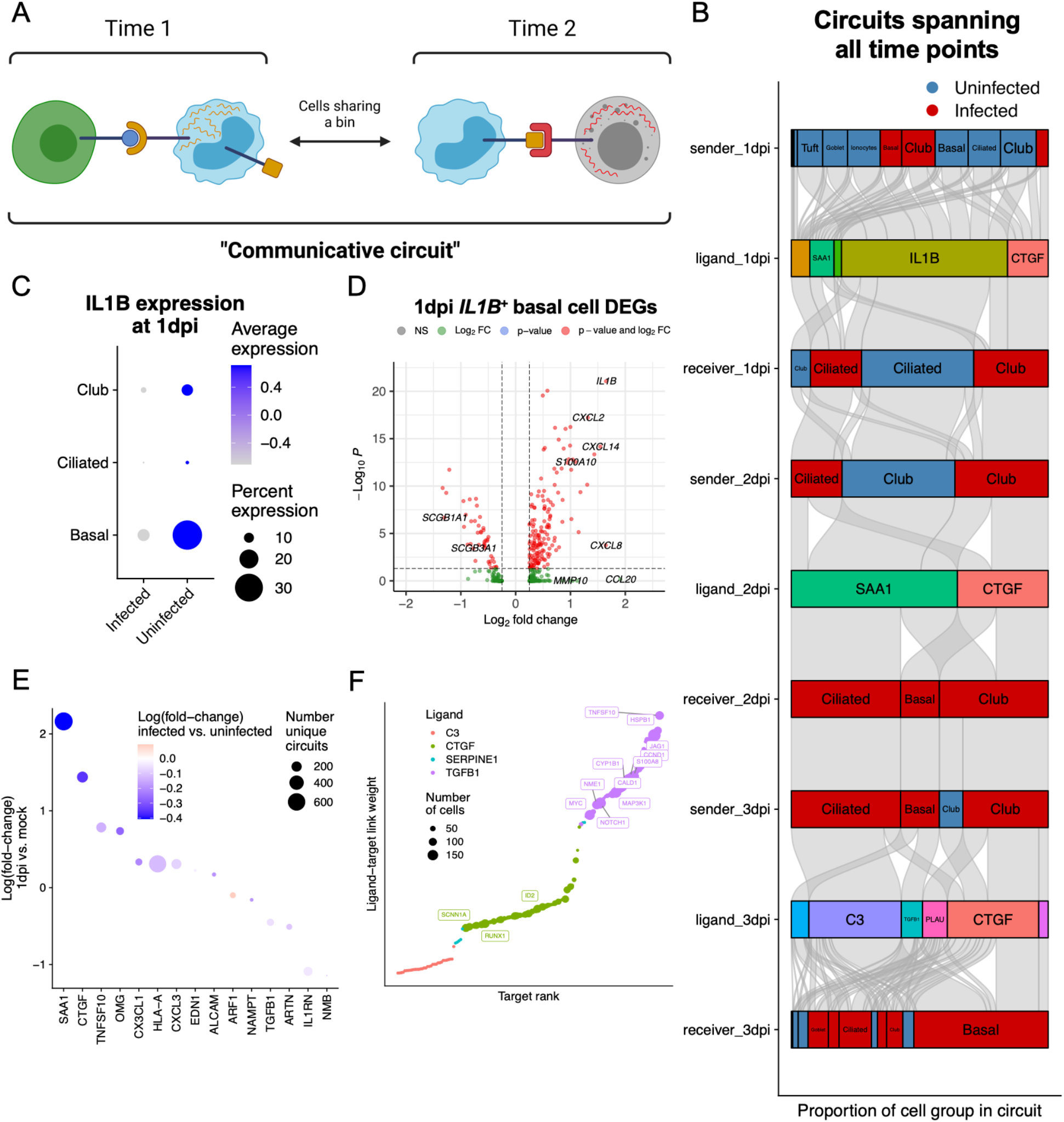
Longitudinal circuits of CCC in acute SARS-CoV-2 infection. **A)** Schematic representing a longitudinal communicative circuit. Four cells participate in a longitudinal circuit if an interaction between S_1_-R_1_ is predicted to result in the upregulation of ligand L_A_ by R_1_, if R_1_ and S_2_ share a bin, and if expression of L_A_ by S_2_ participates in an active communication edge with R_2_. **B)** Alluvial plot depicting longitudinal communicative circuits. Stratum width corresponds to the number of cells in each cell grouping participating in the circuit corrected for the total number of cells in that group. Red strata are infected with SARS-CoV-2; blue strata are composed of uninfected cells. **C)** Dot plot depicting percent and scaled average expression of *IL1B* by club, basal, and ciliated cells at 1 dpi. **D)** Volcano plot depicting log(fold-change) (*x*-axis) and −log(p-value) (*y*-axis) of *IL1B*^+^ basal cells relative to *IL1B*^-^ basal cells at 1 dpi. Positive log(fold-change) indicates the gene is more highly expressed by *IL1B*^+^ basal cells. **E)** Each point represents a ligand predicted to be active and participating in circuits at 1 dpi. The size of each dot represents the number of unique circuits in which that ligand participates. The *y*-axis represents the log(fold-change) of expression of each ligand between 1 dpi (positive *y*-axis) and mock (negative *y-*axis). The color represents the log(fold-change) of expression of each ligand at 1 dpi between infected (red) and uninfected (blue) cells. **F)** Target genes predicted by Scriabin’s implementation of NicheNet^20^ to be upregulated in the receiver cells at the ends of the longitudinal communicative circuits at 3 days post-infection. Points are colored by the active ligand and sized by the number of cells in which the target is predicted to be upregulated by the active ligand.

To illustrate this process, we analyzed a published dataset of SARS-CoV-2 infection in human bronchial epithelial cells (HBECs) in air-liquid interface (ALI) that was sampled daily for 3 days^58^. This dataset contains all canonical epithelial cell types of the human airway and indicates that ciliated and club cells are the preferentially infected cell types in this model system, with some cells having >50% of UMIs from SARS-CoV-2 (**Supplementary FIgure 10**). We first defined a per-cell gene signature of genes variable across time, and used this gene signature to predict active ligands expected to result in the observed cellular gene signatures^20,23^. Next, we used Scriabin’s high-resolution binning workflow to align the datasets from the three post-infection timepoints, which we then used to assemble longitudinal communicative circuits.

Scriabin identified circuits at the level of individual cells that spanned all three post-infection timepoints. We summarized these circuits by author-annotated cell type and whether SARS-CoV-2 reads were detected in the cell (**Figure 5B**). Interestingly, we found that uninfected cells were more frequently the initiators of longitudinal circuits operating over all three time points (**Figure 5B**). The most frequent circuit-initiating ligand was *IL1B* produced by basal, ciliated, and club cells; in these cell types at 1 day post-infection, *IL1B* was more strongly expressed in bystander cells relative to infected cells (**Figure 5C**). Uninfected basal cells at 1 day post-infection displayed the highest expression of *IL1B*, and these *IL1B*^+^ cells were also characterized by higher expression of other pro-inflammatory cytokines including *CCL20* and *CXCL8* (**Figure 5D**). Among the other ligands active at 1 day post-infection, acute phase reactant-encoding genes, including *SAA1* and *CTGF*^*59,60*^, were strongly upregulated at 1 day post-infection relative to the mock condition, and were both more highly expressed by uninfected cells (**Figure 5E**); these genes are known to be induced in the setting of SARS-CoV-2 infection and are hypothesized to be involved in downstream tissue remodeling processes^61^. These results implicate the activity of uninfected bystander cells as potentially important mediators of downstream responses to infection – this may reflect described processes in other viral infections where non-productively infected cells may be key drivers of downstream inflammatory activity^62–64^.

When we assessed the predicted downstream targets at the ends of the longitudinal circuits in both infected and bystander cells, we found that *TGFB1* produced by infected basal cells was predicted to result in the upregulation of *TNFSF10* (encoding TRAIL) and the alarmin *S100A8* predominantly by other infected cells (**Figure 5B**,**F**). Additionally, *TGFB1* was predicted to upregulate both *NOTCH1* and the *NOTCH1* ligand *JAG1*, which indicates that these circuits may induce downstream Notch signaling. In sum, these data illustrate how the single-cell resolution of Scriabin’s CCC analysis workflow can perform integrated longitudinal analyses, nominating hypotheses for experimental validation.

## DISCUSSION

Most existing CCC methodologies aggregate ligand and receptor expression values at the level of the cell type or cluster, potentially obscuring biologically valuable information. Here we introduce a framework to perform comparative analyses of CCC at the level of the individual cell. Scriabin maximally leverages the single-cell resolution of the data to maintain the full structure of both communicative heterogeneity and specificity. We used this framework to find rare communication pathways in the developing intestine known to be required for stem cell maintenance, as well as to define the kinetics of early dynamic communication events in response to SARS-CoV-2 infection through assembly of longitudinal communicative circuits.

A major challenge of single-cell resolved CCC analysis is data inflation: because CCC analysis fundamentally involves performing pairwise calculations on cells or cell groups, it is frequently computationally prohibitive to analyze every sender-receiver cell pair. Importantly, Scriabin implements two complementary workflows to address this issue, both of which avoid the statistically-problematic practices of subsampling and aggregation while maintaining scalability. Subsampling and aggregation preclude a truly comprehensive view of CCC structure and risk obscuring important biology; either can be particularly problematic in situations where a small subset of cells disproportionately drives intercellular communication, with agglomeration potentially concealing the full activity of those cells and subsampling potentially removing those cells altogether. One biological situation in which the preservation of single-cell resolution data could be particularly important is in the setting of activation-induced T cell exhaustion^65^. While exhausted T cells exert divergent effects on their communication targets relative to their activated counterparts, we show that exhausted T cells can often be difficult to distinguish from activated cells by clustering or sub-clustering. By avoiding aggregation and subsampling, Scriabin increases the likelihood of detecting these potentially meaningful differences in CCC pathways.

We observe that aggregation obscures potentially biologically-meaningful subsets of T cells in SCC as well as in reversal reactions in leprosy granulomas. Due to the degree of transcriptional perturbation in T cells during reversal reactions, subclustering is not always a tenable approach to increasing the resolution of CCC analyses because it, in turn, can preclude analysis at a per-sample level. We also show that aggregating across samples, a common practice in existing CCC tools, can return putatively differential CCC edges that are driven disproportionately by individual samples.

As the throughput of scRNA-seq workflows continues to increase, it will be important that single-cell resolution CCC methods are scalable to any dataset size. We introduce two complementary workflows to address this challenge. First, for large comparative analyses, the summarized interaction graph workflow saves computational resources by summarizing the total magnitude of communication between cell-cell pairs and a novel dataset alignment strategy called “binning” enables identification of cells of the greatest biological interest between samples. We provide empirical evidence that this strategy identifies subspaces with the greatest degree of communication perturbation. However, this approach is not robust to situations where ligand-receptor pair mechanisms of CCC change between cell-cell pairs without changing the overall magnitude of CCC.

As an alternative, we also introduce a second single-cell resolution CCC workflow that is scalable to datasets of any size. The interaction program discovery workflow of Scriabin accomplishes this by focusing first on common patterns of ligand-receptor pair co-expression rather than individual cell-cell pairs. Individual cells can be scored for expression of these interaction programs *post hoc*, enabling downstream comparative analyses with a comprehensive view of CCC structure. We apply this workflow to an atlas-scale dataset of human fetal gut development, where we validate a mode of communication between a fibroblast subset and ISCs that has recently been shown to be important for maintaining the ISC niche^51,52^. Due to the relative scarcity of these cells, we show that agglomerative methods fail to discover these important interactions for downstream mechanistic validation.

Longitudinal datasets pose an additional opportunity and challenge for comparative analyses because there is *a priori* knowledge about the sequential relationship between different samples. The single-cell nature of Scriabin’s workflows permits us to analyze how pathways of CCC operate both within and between timepoints in longitudinal datasets. By identifying circuits of CCC that function over multiple timepoints in a longitudinal infection dataset, we can observe how uninfected bystander cells may initiate important inflammatory pathways first which are later amplified by infected cells. A fundamental assumption of the circuit assembly workflow is that ligands upregulated at one timepoint can be observed to exert their biological activity at the following timepoint. This assumption is highly dependent on *a priori* biological knowledge of the communication pathways of interest, as well as on the spacing between timepoints. Assembly of longitudinal communication circuits may represent a valuable strategy to elucidate complex dynamic and temporal signaling events, particularly when longitudinal sampling is performed at frequencies on the same scale as signaling and transcriptional response pathways.

The cell-cell interaction matrix ***M*** is more highly enriched for zero values than gene expression matrices. This is because genes encoding molecules involved in CCC tend to be more lowly expressed than other genes (as the most highly expressed genes tend to encode intracellular proteins involved in cell housekeeping), and because a zero value in *either* the ligand or the receptor of a cell-cell pair will result in a zero value in the interaction vector. Consequently, these zero values can make it difficult for Scriabin to determine if putatively downregulated or “missing” CCC edges are biological or due to dropout. We show that data denoising algorithms for scRNA-seq are capable of removing technical noise caused by data sparsity, substantially improving the yield of bona fide single-cell CCC states. This process can make the presence and absence of CCC edges more interpretable – we recommend the use of denoising algorithms when analyzing datasets generated by low coverage platforms, and particularly for non-unique molecular identifier (UMI) methods which are more likely to be zero-inflated^66,67^.

Another complementary set of techniques for CCC inference are computational methods that infer which cells are communicating by identifying putative multiplets in the dataset, or by directly sequencing interacting cells. The central premise of these techniques, which include Neighbor-seq^68^ and PIC-seq^69^, is that physically interacting cells are likely to be more difficult to dissociate when preparing single-cell suspensions and therefore, that multiplets may be more likely to represent cells that are genuinely interacting. While this provides an additional layer of evidence for biologically-meaningful interactions, there are some communication edges that cannot be detected with these methods. For example, CCC involving secreted ligands will not be adequately modeled with these techniques. Additionally, as each scRNA-seq dataset represents a single snapshot of a sample, cells that have previously interacted but are no longer associated will not be detected. This latter problem has been addressed by techniques such as LIPSTIC^70^ that permanently label cells that have interacted using particular ligands or receptors. However, these methods remain poorly scalable and require prior cell engineering. We anticipate that future technological developments will enable synergy of these complementary approaches towards more comprehensive solutions for CCC analysis.

One current limitation of Scriabin is that it does not take into account situations where multiple receptor subunits encoded by different genes are required in combination to respond to a ligand, or where receptor subunits are known to differentially contribute to collective ligand-receptor avidity. An additional limitation is that Scriabin assumes uniform validity of ligand-receptor interactions in curated protein-protein interaction databases and treats all ligand-receptor pairs as equally important. In situations where it is known *a priori* which ligand-receptor pairs have a higher level of literature support, this information could be used to prioritize downstream analysis of particular ligand-receptor pairs. Additionally, Scriabin assumes the interaction directionality that is presented by the user-selected ligand-receptor database–however, not all interactions are unidirectional and biologically-important receptor-receptor interactions are also possible^71^. Scriabin supports the use of custom ligand-receptor pair databases for users who *a priori* have specific analytical questions involving non-traditional interaction directionality.

Similarly, all downstream signaling analyses in Scriabin rely on NicheNet’s ligand-target activity matrix, which may be biased by the cell types and stimulation conditions used to generate it. The NicheNet database also does not allow for analysis of inhibitory signaling, and thus Scriabin will only return CCC edges predicted to result in activated signaling. While Scriabin uses NicheNet to predict active CCC edges by examining downstream gene expression changes, an additional analysis goal includes identifying the upstream signaling machinery that results in the upregulation of a ligand or denotes successful signaling, as additional power could be gained by using sets of genes to infer upstream signaling rather than relying on ligand expression alone (which could be impacted by dropout or differences between mRNA and protein expression). More generally, Scriabin assumes that gene expression values for ligands and receptors correlate well with their protein expression. A future point of improvement would be to support analysis of multi-modal datasets where cell surface proteins that contribute to CCC are measured directly, or to enable analysis of protein measurements that are imputed from integration with multi-modal references^72^. Future iterations of Scriabin will seek to address these issues, as well as further improve computational efficiency.

Collectively, our work provides a toolkit for comprehensive comparative analysis of CCC in scRNA-seq data, which should empower discovery of information-rich communicative circuitry and niche-phenotype relationships.

## ACKNOWLEDGEMENTS

We thank Drs. William J. Greenleaf and Sam W. Kazer for helpful conversations in the conceptualization of Scriabin’s workflow. We thank Drs. Paul A. Wender and Nancy L-B Weidenbacher for synthesis of Charge-Altering Releasable Transporters. We thank Constantine Tzouanas and Dr. José Ordovas-Montanes for insights on intestinal cell-cell communication pathways. We thank Dr. Andrew Ji for providing processed scRNA-seq data of SCC and matched normals. We also thank all current and former members of the Blish laboratory for helpful discussions of this work. A.J. Wilk is supported by the Stanford Medical Scientist Training Program (T32 GM007365-44) and the Stanford Bio-X Interdisciplinary Graduate Fellowship. Work with Charge-Altering Releasable Transporters was supported by NIH/NIAID R01 AI161803 to C.A.B. This work was also supported by NIH/NIDA DP1 DA04508902 to C.A.B., NIH/NCI 1U54CA217377, U01 28020510, and 1U2CCA23319501 to A.K.S., NIH/NIDA 1DP1DA053731 to A.K.S., Bill and Melinda Gates Foundation INV-027498 and OPP1202327 to A.K.S., the MIT Stem Cell Initiative through Foundation MIT to A.K.S. and a 2019 Sentinel Pilot Project from the Bill and Melinda Gates Foundation to C.A.B./A.K.S..

## AUTHOR CONTRIBUTIONS

A.J.W., A.K.S., S.H., and C.A.B. conceived of the work. A.J.W. built Scriabin, performed computational analyses, cell culture work, flow cytometric analysis, and scRNA-seq profiling. A.J.W. wrote the manuscript with input from all authors. S.H. and C.A.B. jointly supervised the work.

## DATA AVAILABILITY

Raw and processed scRNA-seq data generated in this manuscript will be deposited on GEO without restrictions. All other scRNA-seq data analyzed in this manuscript are publicly available.

## CODE AVAILABILITY

Scriabin is available for download and use as an R package at github.com/BlishLab/scriabin

## METHODS

### Cell-cell interaction matrix workflow

#### Generation of cell-cell interaction matrix

We define the cell-cell interaction vector between a pair of cells as the geometric mean of expression values of each cognate ligand-receptor pair. Formally, the interaction vector *V* between sender cell *N*_*i*_ and receiver cell *N*_*j*_ is given by

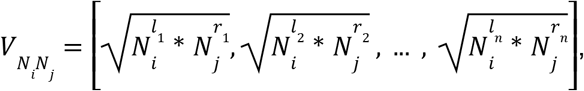

where *l*_*n*_, *r*_*n*_ represent a cognate ligand-receptor pair. We chose to multiply ligand and receptor expression values so that zero values of either ligand or receptor expression would result in a zero value for the corresponding index of the interaction vector. Additionally, we chose to take the square root of the product of ligand-receptor expression values so that highly expressed ligand-receptor pairs do not disproportionately drive downstream analysis. This definition is equivalent to the geometric mean. The cell-cell interaction matrix ***M*** is constructed by concatenating the cell-cell interaction vectors. ***M*** is used as input to low dimensional embeddings for visualization, and nearest neighbor graphs for graph-based clustering.

#### Weighting cell-cell interaction matrix by upstream regulome

The cell-cell interaction matrix ***M*** can be weighted by ligand-receptor edges that are predicted to be active based on observed downstream gene expression changes. First, we identify genes in the dataset that are variable across some axis of interest. For analyses of single datasets, variable genes can be defined as the set of genes with the highest residual variance in the dataset, for example, by calling FindVariableFeatures as implemented by Seurat. For comparative analyses, Scriabin provides several utility functions to aid in the identification of variable genes between samples or between time points, depending on the user’s analytical questions.

Next, the package CelliD^23^, which provides a convenient and scalable workflow to define single-cell gene signatures, is used to define per-cell gene signatures. Briefly, user-defined variable genes are used to embed the dataset into low dimensional space by multiple correspondence analysis (MCA). A cell’s gene signature is then defined as the set of genes to which that cell is nearest in the MCA biplot. A quantile cutoff is used to threshold gene proximity, by default the 5% of nearest genes.

NicheNet’s^20^ ligand-target matrix, which denotes the regulatory potential scores between ligands and target genes, is then used to rank ligands based on their predicted ability to result in the per-cell gene signature. First, expressed genes are defined by the percentage of cells in which they are detected (by default, 2.5%). Next, a set of potential ligands is defined as those ligands which are expressed genes and for which at least one receptor is also an expressed gene.

Next, the ligand-target matrix is filtered to contain only the set of potential ligands and targets in the set of expressed genes. The authors of NicheNet have shown that the Pearson correlation coefficient between a ligand’s target prediction and observed transcriptional response is the most informative metric of ligand activity^20^. Therefore, the activity of a single ligand for a single cell is defined as the Pearson correlation coefficient between the vector of that cell’s gene signature and the target gene scores for that ligand. For each active ligand, target gene weights for each cell are defined as the ligand-target matrix regulatory score for the top 250 targets for each ligand that appear in a given cell’s gene signature. We select a Pearson coefficient threshold (by default 0.075) to define active ligands in each cell.

Finally, we weight individual values of 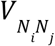. Scriabin supports two methods for weighting the CCIM by predicted ligand activities. Method “product” (default) weights interaction vectors proportionally to predicted ligand activities. The vector of ligand activities for receiver cell *N*_*j*_, *A*_*i*_, is scaled so that values above the pearson threshold lie between two scaling factors (by default 1.5 and 3), and values below the pearson threshold are set to 1. The interaction vector is then given by:

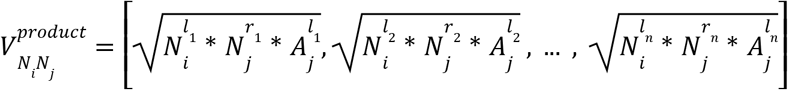

Method “sum” treats a ligand activity as orthogonal evidence of receptor expression. Pearson coefficients in the vector of ligand activities for receiver cell *N*_*j*_, *A*_*j*_, that are below the Pearson threshold are set to 0. The interaction vector is then given by:

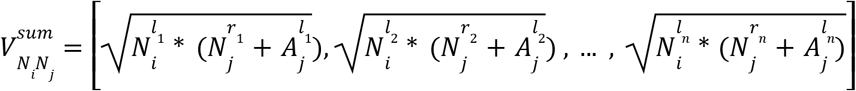

Use cases for ligand activity weighting methods, as well as other parameters involved in calculating ligand activities, are described in the **Supplementary Text**.

#### Downstream analysis of weighted cell-cell interaction matrices

***M*** can be treated analogously to the gene expression matrix and used for downstream analysis tasks like dimensionality reduction. After generation and (optional) weighting of ***M*** by active ligands, ***M*** is placed into an assay of a Seurat object for downstream analysis. ***M*** is scaled by ScaleData, latent variables found via PCA, and the top principal components (identified by ElbowPlot for each dataset) used to embed the dataset in two dimensions using UMAP^73^. Neighbor graphs are constructed by FindNeighbors, which can then be clustered via modularity optimization graph-based clustering^74^ as implemented by Seurat’s FindClusters^72^. Differential ligand-receptor edges between clusters, cell types, or samples can be identified via FindMarkers. Scriabin provides several utility functions to facilitate visualization of gene expression profiles or other metadata on Seurat objects built from cell-cell interaction matrices.

### Summarized interaction graph and binning workflow

#### Generation of summarized interaction graph

Because ***M*** scales exponentially with dataset size, it is frequently impractical to calculate ***M*** for all cell-cell pairs *N*_*i*_,*N*_*j*_. In this situation, Scriabin supports two workflows that do not require aggregation or subsampling. In the first workflow, a summarized cell-cell interaction graph ***S*** is built in lieu of ***M*** where 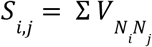. ***S*** thus represents the magnitude of predicted interaction across all cognate ligand-receptor pairs expressed by all sender-receiver cell pairs. ***S*** is then corrected for associations with sequencing depth by linear regression. The sequencing depth of cell-cell pair *N*_*i*_,*N*_*j*_ is defined as 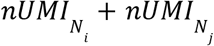. A linear model is fit to describe the relationship between the summarized interaction score (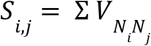, where *S* is the summarized interaction matrix and 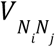 is the interaction vector for cell-cell pair *N*_*i*_,*N*_*j*_) and the total sequencing depth of each cell-cell pair. The residuals of this model are used as a sequencing depth-corrected ***S. S*** may optionally be weighted through prediction of ligand activity as described above. The second workflow is described below under “**Interaction program discovery workflow**”.

#### Dataset binning for comparative CCC analyses

Once summarized interaction graphs are built for multiple samples, alignment of these graphs requires knowledge about which cells between samples represent a shared molecular state. The goal of binning is to assign each cell a bin identity so that ***S*** from multiple samples can be summarized into equidimensional matrices based on shared bin identities.

The binning process begins by constructing a shared nearest neighbor (SNN) graph via FindNeighbors defining connectivity between all cells to be compared. Alternate neighbor graphs, for example those produced using Seurat’s weighted nearest neighbor workflow which leverage information from multi-modal references, can also be used. Next, mutual nearest neighbors (MNNs) are identified between all sub-datasets to be compared via Seurat’s integration workflow (FindIntegrationAnchors)^21^. Briefly, two sub-datasets to be compared are placed into a shared low dimensional space via diagonalized canonical correlation analysis (CCA), and the canonical correlation vectors are log-normalized. Normalized canonical correlation vectors are then used to identify k-nearest neighbors for each cell in its paired dataset and the resulting MNN pairings are scored as described^21^. Low scoring MNN pairings are then removed, as they have a higher tendency to represent incorrect cell-cell correspondences when orthogonal data is available (**Supplementary Figure 6**).

For each cell that participates in an MNN pair, Scriabin defines a bin as that cell and all cells with which it participates in an MNN pair. Considering a dataset of cells ***i*** of which a subset ***i’*** participate in an MNN pair, for each cell ***i’***_***n***_ we define a bin ***j***_***n***_ which contains ***i’***_***n***_ and all MNNs of ***i’***_***n***_. Next, Scriabin constructs a connectivity matrix ***G*** where ***G***_***i***,***j***_ is the mean connectivity in the SNN graph between cell ***i*** and the cells within bin ***j***. Each cell ***i***_***n***_ is assigned a bin identity of the bin ***j***_***n***_ with which it shares the highest connectivity in ***G***. Thus, at the end of this process each cell has a single bin identity which reflects its SNN similarity to a group of cells with cross-dataset MNN connectivity.

However, at this stage each bin ***j***_***n***_ may not contain cells from all the samples being compared. Thus, we next optimize for the set of bins that results in the best representation of all samples. Bins ***j*** with the lowest total connectivity and lowest multi-sample representation in ***G*** are iteratively removed and cell bin identities re-scored until the mean sample representation of each bin plateaus. Within-bin connectivity and sample representation are further improved by re-assigning cells that result in better sample representation of an incompletely represented bin while maintaining equal or greater SNN connectivity with the cells in that bin. Finally, remaining incompletely represented bins are merged with the nearest completely represented bin with which it shares the highest SNN connectivity. At the end of this process, each cell will thus have a single assigned bin identity, where each bin contains cells from all samples to be compared.

#### Statistical analysis of bin significance

Bins are then tested for the statistical significance of their connectivity structure using a permutation test. For each bin, random bins of the same size and number of cells per sample are generated iteratively (by default 10,000 times). The connectivity vector of the real bins is tested against each of the random bins by a one-sided Mann-Whitney U test. If the bin fails 500 or more of these tests (p-value 0.05), it is considered non-significant.

Because bin SNN connectivity is generally non-zero, but randomly sampled cells generally have an SNN connectivity of zero, this strategy will tend to return most bins as statistically significantly connected. Thus, we recommend passing high-resolution cell type labels to the binning significance testing. In this situation, randomly generated bins are generated by randomly selecting cells from the same sample and cell type annotation, and the permutation test proceeds as described above. Bins where greater than a threshold (by default 95%) of cells belong to the same cell type annotation are automatically considered significant. This avoids rare cell types that may only form a single bin from being discarded. Cells that were assigned to bins which failed the significance testing are re-assigned to the bin with which they share the highest SNN connectivity.

#### Identification of variable bins

For each bin, a Kruskal-Wallis test is used to assess differences in the magnitude of CCC between cell-cell pairs from different samples. The Kruskal-Wallis p-value and test statistic can be used to identify which bins contain cells that exhibit the highest change in prediction interaction scores. Specific samples that contribute to each significantly variable bin’s perturbation are then identified through Dunn’s post-hoc test. This set of sender and receiver cells can then be used to construct ***M*** as described above.

### Interaction program discovery workflow

#### Iterative approximation of a ligand-receptor pair topological overlap matrix (TOM)

An alternative to the summarized interaction graph workflow is to instead identify co-expressed ligand-receptor pairs, which we refer to as “interaction programs.” This approach represents an adaptation of the well-established weighted gene correlation network analysis (WGCNA)^22^ and is scalable to any dataset size and still permits analysis of CCC at single-cell resolution. The first step in this workflow is to generate a signed covariance matrix of ligand-receptor pairs for each sample, defined as:

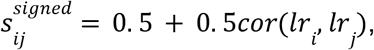

where *lr*_*i*_, *lr*_*j*_ are individual ligand-receptor pair vectors of ***M***. In large datasets, 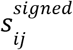 is approximated by iteratively generating subsets of ***M***. 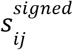 is next converted into an adjacency matrix via soft thresholding:

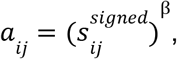

where β is the soft power. Soft power is a user-defined parameter that is recommended to be the lowest value that results in a scale-free topology model fit of > 0.8. Next, this adjacency matrix is converted into a TOM as described^75^. This process proceeds separately for each sample to be analyzed in a multi-sample dataset.

#### Identification and significance testing of interaction programs

The TOM is hierarchically clustered, and interaction programs identified through adaptive branch pruning of the hierarchical clustering dendrogram. Intramodular connectivity for each ligand-receptor pair in each interaction program is then calculated as described^76^. If interaction programs are being discovered in a multi-sample dataset, similar modules (defined by Jaccard overlap index above a user-defined threshold) are merged. Next, interaction programs are then tested for statistically significant co-expression structure via a permutation test where random interaction programs are generated 10,000 times. The correlation vector of the real module is tested against each of the random modules by a one-sided Mann-Whitney U test. If the module fails 500 or more of these tests (p-value 0.05), it is considered non-significant. Each sample is tested for significant correlation of each module.

#### Downstream analysis of interaction programs

Single cells are scored separately for the expression of the ligands and receptors of each significant module with Seurat’s AddModuleScore. This function calculates a module score by comparing the expression level of an individual query gene to other randomly-selected control genes expressed at similar levels to the query genes, and is therefore robust to scoring modules containing both lowly and highly expressed genes, as well as to scoring cells with different sequencing depth. Scriabin includes several utility functions to conveniently visualize interaction program expression for sender and receiver cells.

### Identification of longitudinal CCC circuits

A longitudinal CCC circuit is composed of S_1_-L_1_-R_1_-S_2_-L_2_-R_2_, where S are sender cells and R are receiver cells at timepoints 1 and 2, and where L_1_ is expressed by/sensed by S_1_/R_1_ and L_2_ is expressed by/sensed by S_2_/R_2_. For computational efficiency, construction of longitudinal CCC circuits starts at the end of the circuit and proceeds upstream. First, ligands L_2_ predicted by NicheNet to be active in receiver cells at timepoint 2 are identified. Next, sender cells that express L_2_ *and* have the L_2_ in its per-cell gene signature are identified. Among the bins occupied by these S_2_ candidates, Scriabin then searches for receiver cells at timepoint 1 that occupy the same bin and have the corresponding timepoint 2 ligand L_2_ within its list of upregulated target genes and identifies the ligand(s) L_1_ predicted by NicheNet to result upregulation of that target. Finally, Scriabin identifies S_1_ candidates that express the timepoint 1 ligands L_1_ and have L_1_ in its per-cell gene signature. S_1_-R_1_-S_2_-R_2_ cell groups that meet these criteria are retained for further analysis. This process repeats for every pair of timepoints. Finally, Scriabin searches for overlap between circuits of sequential time point pairs to identify circuits that operate over more than two timepoints.

### Ligand-receptor pair databases for analysis

Scriabin supports the use of 15 ligand-receptor interaction databases for all analytical functions; these resources were collected from Liana^77^. By default, Scriabin uses the OmniPath database^18,19^ filtered for curation strength of > 7 to ensure that ligand-receptor interactions with strong experimental evidence are included in downstream analysis. Scriabin also supports the use of custom ligand-receptor pair lists for users with specific analytical questions.

### Transfection and co-culture of primary NK and B cells

Peripheral blood mononuclear cells (PBMCs) were acquired from a healthy blood donor that was consented for release of genetic data by the Stanford Blood Center. PBMCs were isolated by Ficoll-Paque (GE Healthcare) density gradient centrifugation and cryopreserved in 90% FBS + 10% DMSO (v/v). PBMCs were thawed at 37°C in complete RPMI-1640 media (supplemented with 10% FBS, L-glutamine, and Penicillin-Streptomycin-Amphotericin; RP10) containing benzonase (EMD Millipore). NK cells and B cells were purified from thawed PBMCs by magnetic bead isolation *via* negative selection according to the manufacturer’s specifications (Miltenyi, cat. 130-092-657 and 130-101-638, respectively). NK and B cells were maintained in complete RP10 media without additional cytokines to ensure a resting state. All cell culture was performed at 37°C/5% CO_2_ in a humidified environment.

eGFP-, CD40-, and CD40L-encoding mRNAs were purchased from TriLink Biotech and used without further purification. Notably, open-reading frame (ORF) sequences for mRNAs encoding CD40 and CD40L were codon optimized using the codon optimization tool developed by Integrated DNA Technologies: this serves both to improve translational efficacy as well as to enable distinguishing endogenous vs. exogenous *CD40* and *CD40LG* mRNAs through sequencing.

mRNAs were delivered to isolated NK and B cells via transfection by Charge-Altering Releasable Transporters (CARTs) as previously described^78^. Briefly, CART/mRNA polyplexes were prepared by diluting 0.84 of mRNA (1 μg/μl) into 14.52 μL PBS (pH 5.5). To this solution was added 1.44 μL of CART BDK-O_7_:N_7_:A_13_ (2 mM DMSO) to achieve a charge ratio of 10:1 (+/-, assuming all ionizable cationic groups are protonated). After mixing by finger vortex for 15 seconds, 2.5 μL of the polyplexes were added to cells and incubated for 6 h in serum-free media. Following this incubation, an aliquot was taken from each transfection condition for flow cytometric analysis, FBS was added to a final concentration of 10%, the cells counted, and NK cells and B cells from the respective transfection conditions mixed together in a 1:1 ratio for co-culture. Cells were co-cultured for 12 hours before analysis by flow cytometry and scRNA-seq.

### Flow cytometry

Antibodies used for flow cytometric analyses are listed in **Supplementary Table 2**. eBioscience™ Fixable Viability Dye eFluor™ 780 (ThermoFisher) was used as a viability stain. Following application of viability stain, cells were surface stained for 20 minutes at room temperature, before acquisition on an Aurora flow cytometer (Cytek Biosciences) and analysis by FlowJo version 10.6.1.

### scRNA-seq by Seq-Well

The Seq-Well platform for scRNA-seq was utilized as described previously^56,79–82^. Immediately following co-culture, cells were counted and diluted in RP10 to a concentration of 75,000 cells/mL. 200 μL of this cell suspension (15,000 cells) was then loaded onto Seq-Well arrays pre-loaded with mRNA capture beads (ChemGenes). Following four washes with DPBS to remove serum, the arrays were sealed with a polycarbonate membrane (pore size of 0.01 µm) for 30 minutes at 37°C. Next, arrays were placed in lysis buffer, transcripts hybridized to the mRNA capture beads, and beads recovered from the arrays and pooled for downstream processing. Immediately after bead recovery, mRNA transcripts were reverse transcribed using Maxima H-RT (Thermo Fisher EPO0753) in a template-switching-based RACE reaction, excess unhybridized bead-conjugated oligonucleotides removed with Exonuclease I (NEB M0293L), and second-strand synthesis performed with Klenow fragment (NEB M0212L) to enhance transcript recovery in the event of failed template switching^80^. Whole transcriptome amplification (WTA) was performed with KAPA HiFi PCR Mastermix (Kapa Biosystems KK2602) using approximately 6,000 beads per 50 μL reaction volume. Resulting libraries were then pooled in sets of 6 (approximately 36,000 beads per pool) and products purified by Agencourt AMPure XP beads (Beckman Coulter, A63881) with a 0.6x volume wash followed by a 0.8x volume wash. Quality and concentration of WTA products was determined using an Agilent TapeStation, with a mean product size of >800bp and a non-existent primer peak indicating successful preparation. Library preparation was performed with a Nextera XT DNA library preparation kit (Illumina FC-131-1096) with 1 ng of pooled library using single-index primers. Tagmented and amplified libraries were again purified by Agencourt AMPure XP beads with a 0.6x volume wash followed by a 1.0x volume wash, and quality and concentration determined by TapeStation analysis. Libraries between 400-1000bp with no primer peaks were considered successful and pooled for sequencing. Sequencing was performed on a NovaSeq 6000 instrument (Illumina; Chan Zuckerberg Biohub). The read structure was paired-end with read 1 beginning from a custom read 1 primer^79^ containing a 12bp cell barcode and an 8 bp unique molecular identifier (UMI), and with read 2 containing 50bp of mRNA sequence.

### Alignment and quality control of scRNA-seq data

Sequencing reads were aligned and count matrices assembled using STAR^83^ and dropEst^84^, respectively. Briefly, the mRNA reads in read 2 demultiplexed FASTQ files were tagged with the cell barcode and UMI for the corresponding read in the read 1 FASTQ file using the dropTag function of dropEst. Next, reads were aligned with STAR using the GRCh38.p13 (hg38) human reference genome from Ensembl. This reference also included sequences and annotations for the codon-optimized ORFs for GFP-, CD40-, and CD40L-encoding mRNAs so that both endogenous and exogenous mRNAs could be quantified. Count matrices were built from resulting BAM files using dropEst^84^. Cells that had fewer than 750 UMIs or greater than 15,000 UMIs, as well as cells that contained greater than 20% of reads from mitochondrial genes or rRNA genes (*RNA18S5* or *RNA28S5*), were considered low quality and removed from further analysis. To remove putative multiplets, cells that expressed more than 75 genes per 100 UMIs were also filtered out.

### Pre-processing of scRNA-seq data

The R package Seurat^21,72,85^ was used for data scaling, transformation, clustering, dimensionality reduction, differential expression analysis, and most visualizations. Unless otherwise noted, data were scaled and transformed and variable genes identified using the SCTransform() function, and linear regression performed to remove unwanted variation due to cell quality (% mitochondrial reads, % rRNA reads). PCA was performed using the 3,000 most highly variable genes, and the first 50 principal components (PCs) used to perform UMAP to embed the dataset into two dimensions^73,86^. Next, the first 50 PCs were used to construct a shared nearest neighbor graph (SNN; FindNeighbors()) and this SNN used to cluster the dataset (FindClusters()). Although upstream quality control removed many dead or low quality cells, if any clusters were identified that were defined by few canonical cell lineage markers and enriched for genes of mitochondrial or ribosomal origin, these clusters were removed from further analysis^87,88^.

### Annotation of transfected NK and B cells in scRNA-seq data

Because of the strong degree of transcriptional perturbation caused by transfection (**Supplementary Figure 1**), we elected to annotate NK and B cells in this dataset by integration with a multimodal reference rather than by graph-based clustering. First, we noted two clusters with high expression of CD3-encoding genes and monocyte-specific genes (including *LYZ* and *CD14*), respectively; we considered these clusters contaminating T cells and monocytes and removed them from further analysis. Next, we used the multimodal (whole transcriptome plus 228 cell surface proteins) PBMC dataset published by Hao, et al.^72^ as a reference. We subsetted the reference to contain only NK and B cells, scaled both the transcriptome and protein assays, and ran PCA on both modalities. Next, we found multimodal neighbors between the modalities via weighted nearest neighbor (WNN) analysis, which learns the relative utility of each data modality in each cell. Supervised PCA (SPCA) was then run on the WNN SNN graph, which seeks to capture a linear transformation that maximizes its dependency to the WNN SNN graph. These SPCA reduced dimensions were then used for identification of anchors between the reference and query datasets as previously described^21^. Finally, these anchors were used to transfer reference cell type annotations to the query dataset.

### Processing, analysis, and visualization of public scRNA-seq datasets

Datasets of PBMCs (pbmc5k and pbmc10k) were downloaded from 10X genomics; for comparison of mouse and human PBMCs, datasets from 10X Genomic’s cell multiplexing oligo demonstration were used (https://www.10xgenomics.com/resources/datasets). Processed scRNA-seq data of SCC and matched normals^29^ were provided directly by the study authors.

Processed CRISPRa Perturb-seq data were downloaded from Zenodo record #5784651^40^. scRNA-seq data of human leprosy granulomas^41^ were downloaded from https://github.com/mafeiyang/leprosy_amg_network. Data from developing fetal intestine^49^ were acquired from the cellxgene portal: https://cellxgene.cziscience.com/collections/60358420-6055-411d-ba4f-e8ac80682a2e. Data of longitudinal responses to SARS-CoV-2 infection in HBECs^58^ were downloaded from the Gene Expression Omnibus accession #GSE166766. In each case, we acquired raw count matrices or processed Seurat objects containing raw count matrices. Any upstream processing was performed as described in the respective manuscripts.

Raw count matrices from Ravindra, et al.^58^ required filtering before downstream analysis; cells meeting the following criteria were kept: >1,000 UMIs, <20,000 UMIs, >500 unique features, <0.85 UMI-to-unique feature ratio, <20% UMIs of mitochondrial origin, <35% reads from ribosomal protein-encoding genes. Pbmc5k and pbmc10k datasets from 10X genomics were filtered to enforce a minimum number features per cell of 200 and to remove genes not expressed in at least 3 cells.

Cell type annotations were provided for the Ji, et al., Ma, et al., and Fawkner-Corbett, et al. datasets, which were used for downstream analytical tasks. For the Ravindra, et al. dataset, manual annotation of cellular identity was performed by finding differentially expressed genes for each cluster using Seurat’s implementation of the Wilcoxon rank-sum test (FindMarkers()) and comparing those markers to known cell type-specific genes listed in the Ravindra, et al.^58^ PBMC datasets were annotated by weighted nearest neighbor projection and label transfer from a multi-modal PBMC reference as described^72,82^.

For analysis of T cell exhaustion in the SCC dataset from Ji, et al.^29^, an exhaustion signature was defined by: *PDCD1, TOX, CXCL13, CTLA4, TNFRSF9, HAVCR2, LAG3, CD160*, and *CD244*. This signature incorporates several markers of exhausted T cell reported in the literature^65,89–91^. Individual T cells were scored for expression of this signature using Seurat’s AddModuleScore.

### Comparative analyses between Scriabin and published CCC analysis methods

Pbmc5k and pbmc10k datasets from 10X genomics were used to benchmark the computational efficiency of Scriabin. For single dataset analyses, pbmc5k was randomly subsetted to multiple dataset sizes. Cell type annotations were passed to Connectome^38^, NATMI^17^, CellChat^15^, iTALK^16^, and SingleCellSignalR (SCA)^37^, which were run using default parameters defined by Liana^39^. The time for these methods to return results was compared to a version of Scriabin that generated and visualized a full-dataset summarized interaction graph and returned pseudobulk ligand-receptor pair scores for each cell type annotation. Connectome^38^ is the only of these packages that supports a full comparative workflow. For comparative analysis, we analyzed differences in CCC between the pbmc5k and pbmc10k datasets. We compared Connectome’s total runtime to the runtime of Scriabin to generate full dataset summarized interaction graphs, perform dataset binning, and visualize the most perturbed bins.

Multiple ligand-receptor resources compiled by Liana^39^ were used to compare results returned by published CCC analysis methods and Scriabin. The following results parameters were used from each method: prob (CellChat), LRscore (SingleCellSignalR), weight_norm (Connectome), weight_comb (iTALK), edge_avg_expr (NATMI). To visualize the overlap in results between the methods and resources, we extracted the top 1,000 results from each method-resource pair and calculated the Jaccard index between these top results (as described by ^39^).

### Analysis of spatial transcriptomic datasets with Scriabin

To evaluate if Scriabin returns biologically meaningful CCC edges, we downloaded spatial coordinates and gene expression count matrices from 10 spatial transcriptomic datasets from the 10X Visium platform available at https://www.10xgenomics.com/resources/datasets. We also analyzed a spatial transcriptomic dataset published by Ma, et al. of a human leprosy granuloma^41^. We treated each count matrix analogously to scRNA-seq data, performing data transformation and dimensionality reduction as described above. We calculated per-cell gene signatures for each dataset based on variable genes across the dataset, which we then used to rank ligands based on their predicted ability to result in the observed gene expression profile using NicheNet^20^. Next, we constructed a summarized interaction graph using a ligand-receptor pair database that was restricted to membrane-bound ligands and receptors, which we weighted according to the predicted ligand activities. Finally, we compared the distance quantile of the top 1% of interacting cell-cell pairs compared to randomly permuted distances.

### Analysis of CRISPRa Perturb-seq data

To quantify Scriabin’s ability to detect changes in CCC at single-cell resolution, we analyzed data from a pooled genetic perturbation screen. We elected to analyze the Perturb-seq dataset published by Schmidt, et al.^40^ as this dataset was collected on primary cells and contained a high number of gRNAs (15) targeting cell surface ligands or receptors used in CCC. We collected a processed and publicly available h5Seurat object of the anti-human CD3/CD28 restimulated T cells from the Schmidt, et al. dataset from zenodo.org/record/5784651. The authors’ gRNA calls were used for all downstream analysis; we identified gRNAs ***g*** in this dataset that targeted cell-surface ligands or receptors that were present in OmniPath’s ligand-receptor interaction database. The dataset was then subsetted to include only cells transduced with a gRNA targeting one of these cell-surface ligands or receptors, or cells trasudced with a non-targeting gRNA. Untransduced T cells were removed from further analysis. We repeated the following process for each gRNA ***g***_***i***_. Given a gRNA, ***g***_***A***_, targeting a ligand-encoding gene ***A***: we isolated cells transduced with ***g***_***A***_ and cells transduced with a non-targeting gRNA. From this subsetted dataset we generated a CCIM without ligand activity ranking using Scriabin’s CCIM workflow. We next isolated interaction vectors ***V*** for ligand ***A*** and all receptors of ***A, R***_***A***_. For each interaction vector ***V***_***AR***_, we constructed an ROC curve using ***V***_***AR***_ as the predictor variable and the gRNA assignment (either ***g***_***A***_ or non-targeting) as the response variable to quantify and visualize the sensitivity and specificity of the prediction.

## REFERENCES

1. Almet, A. A., Cang, Z., Jin, S. & Nie, Q. The landscape of cell–cell communication through single-cell transcriptomics. Current Opinion in Systems Biology 26, 12–23 (2021).

2. Armingol, E., Officer, A., Harismendy, O. & Lewis, N. E. Deciphering cell–cell interactions and communication from gene expression. Nat. Rev. Genet. 22, 71–88 (2020).

3. Tanay, A. & Regev, A. Scaling single-cell genomics from phenomenology to mechanism. Nature 541, 331–338 (2017).

4. Yosef, N. & Regev, A. Writ large: Genomic dissection of the effect of cellular environment on immune response. Science (2016) doi:10.1126/science.aaf5453.

5. Ramilowski, J. A. et al. A draft network of ligand–receptor-mediated multicellular signalling in human. Nat. Commun. 6, 7866 (2015).

6. Dixit, A. et al. Perturb-Seq: Dissecting Molecular Circuits with Scalable Single-Cell RNA Profiling of Pooled Genetic Screens. Cell 167, 1853–1866.e17 (2016).

7. Schraivogel, D. et al. Targeted Perturb-seq enables genome-scale genetic screens in single cells. Nat. Methods 17, 629–635 (2020).

8. Puram, S. V. et al. Single-Cell Transcriptomic Analysis of Primary and Metastatic Tumor Ecosystems in Head and Neck Cancer. Cell 171, 1611–1624.e24 (2017).

9. Camp, J. G. et al. Multilineage communication regulates human liver bud development from pluripotency. Nature 546, 533–538 (2017).

10. Pavličev, M. et al. Single-cell transcriptomics of the human placenta: inferring the cell communication network of the maternal-fetal interface. Genome Res. 27, 349–361 (2017).

11. Zepp, J. A. et al. Distinct Mesenchymal Lineages and Niches Promote Epithelial Self-Renewal and Myofibrogenesis in the Lung. Cell 170, 1134–1148.e10 (2017).

12. Cohen, M. et al. Lung Single-Cell Signaling Interaction Map Reveals Basophil Role in Macrophage Imprinting. Cell 175, 1031–1044.e18 (2018).

13. Vento-Tormo, R. et al. Single-cell reconstruction of the early maternal-fetal interface in humans. Nature 563, 347–353 (2018).

14. Raredon, M. S. B. et al. Single-cell connectomic analysis of adult mammalian lungs. Sci Adv 5, eaaw3851 (2019).

15. Jin, S. et al. Inference and analysis of cell-cell communication using CellChat. Nat. Commun. 12, 1–20 (2021).

16. Wang, Y. et al. iTALK: an R Package to Characterize and Illustrate Intercellular Communication. doi:10.1101/507871.

17. Hou, R., Denisenko, E., Ong, H. T., Ramilowski, J. A. & Forrest, A. R. R. Predicting cell-to-cell communication networks using NATMI. Nat. Commun. 11, 5011 (2020).

18. Türei, D., Korcsmáros, T. & Saez-Rodriguez, J. OmniPath: guidelines and gateway for literature-curated signaling pathway resources. Nat. Methods 13, 966–967 (2016).

19. Türei, D. et al. Integrated intra- and intercellular signaling knowledge for multicellular omics analysis. Mol. Syst. Biol. 17, e9923 (2021).

20. Browaeys, R., Saelens, W. & Saeys, Y. NicheNet: modeling intercellular communication by linking ligands to target genes. Nat. Methods 17, 159–162 (2020).

21. Stuart, T. et al. Comprehensive Integration of Single-Cell Data. Cell 177, 1888–1902.e21 (2019).

22. Langfelder, P. & Horvath, S. WGCNA: an R package for weighted correlation network analysis. BMC Bioinformatics 9, 559 (2008).

23. Cortal, A., Martignetti, L., Six, E. & Rausell, A. Gene signature extraction and cell identity recognition at the single-cell level with Cell-ID. Nat. Biotechnol. 39, 1095–1102 (2021).

24. McMurdie, P. J. & Holmes, S. Waste not, want not: why rarefying microbiome data is inadmissible. PLoS Comput. Biol. 10, e1003531 (2014).

25. Haghverdi, L., Lun, A. T. L., Morgan, M. D. & Marioni, J. C. Batch effects in single-cell RNA-sequencing data are corrected by matching mutual nearest neighbors. Nat. Biotechnol. 36, 421–427 (2018).

26. Wang, J. et al. Evaluation of ultra-low input RNA sequencing for the study of human T cell transcriptome. Sci. Rep. 9, 1–13 (2019).

27. Wu, L. et al. Blockade of TIGIT/CD155 Signaling Reverses T-cell Exhaustion and Enhances Antitumor Capability in Head and Neck Squamous Cell Carcinoma. Cancer Immunol Res 7, 1700–1713 (2019).

28. Kiner, E. et al. Gut CD4+ T cell phenotypes are a continuum molded by microbes, not by TH archetypes. Nat. Immunol. 22, 216–228 (2021).

29. Ji, A. L. et al. Multimodal Analysis of Composition and Spatial Architecture in Human Squamous Cell Carcinoma. Cell 182, 497–514.e22 (2020).

30. Joller, N. & Kuchroo, V. K. Tim-3, Lag-3, and TIGIT. Curr. Top. Microbiol. Immunol. 410, 127–156 (2017).

31. van Dijk, D. et al. Recovering Gene Interactions from Single-Cell Data Using Data Diffusion. Cell 174, 716–729.e27 (2018).

32. Linderman, G. C. et al. Zero-preserving imputation of single-cell RNA-seq data. Nat. Commun. 13, 1–11 (2022).

33. Lawlor, N. et al. Single-cell transcriptomes identify human islet cell signatures and reveal cell-type-specific expression changes in type 2 diabetes. Genome Res. 27, 208–222 (2017).

34. Baron, M. et al. A Single-Cell Transcriptomic Map of the Human and Mouse Pancreas Reveals Inter- and Intra-cell Population Structure. Cell Syst 3, 346–360.e4 (2016).

35. Zilionis, R. et al. Single-cell barcoding and sequencing using droplet microfluidics. Nat. Protoc. 12, 44–73 (2016).

36. Korsunsky, I. et al. Fast, sensitive and accurate integration of single-cell data with Harmony. Nat. Methods 16, 1289–1296 (2019).

37. Cabello-Aguilar, S. et al. SingleCellSignalR: inference of intercellular networks from single-cell transcriptomics. Nucleic Acids Res. 48, e55–e55 (2020).

38. Raredon, M. S. B. et al. Connectome: computation and visualization of cell-cell signaling topologies in single-cell systems data. bioRxiv 2021.01.21.427529 (2021) doi:10.1101/2021.01.21.427529.

39. Dimitrov, D. et al. Comparison of Resources and Methods to infer Cell-Cell Communication from Single-cell RNA Data. bioRxiv 2021.05.21.445160 (2021) doi:10.1101/2021.05.21.445160.

40. Schmidt, R. et al. CRISPR activation and interference screens decode stimulation responses in primary human T cells. Science 375, eabj4008 (2022).

41. Ma, F. et al. The cellular architecture of the antimicrobial response network in human leprosy granulomas. Nat. Immunol. 22, 839–850 (2021).

42. Gordon, S. Alternative activation of macrophages. Nat. Rev. Immunol. 3, 23–35 (2003).

43. Ridley, D. S. & Jopling, W. H. Classification of leprosy according to immunity. A five-group system. Int. J. Lepr. Other Mycobact. Dis. 34, 255–273 (1966).

44. Flynn, J. L. et al. An essential role for interferon gamma in resistance to Mycobacterium tuberculosis infection. J. Exp. Med. 178, 2249–2254 (1993).

45. Herbst, S., Schaible, U. E. & Schneider, B. E. Interferon gamma activated macrophages kill mycobacteria by nitric oxide induced apoptosis. PLoS One 6, e19105 (2011).

46. Ní Cheallaigh, C. et al. A Common Variant in the Adaptor Mal Regulates Interferon Gamma Signaling. Immunity 44, 368–379 (2016).

47. Verhagen, C. E. et al. Reversal reaction in borderline leprosy is associated with a polarized shift to type 1-like Mycobacterium leprae T cell reactivity in lesional skin: a follow-up study. J. Immunol. 159, 4474–4483 (1997).

48. Teles, R. M. B. et al. Identification of a systemic interferon-γ inducible antimicrobial gene signature in leprosy patients undergoing reversal reaction. PLoS Negl. Trop. Dis. 13, e0007764 (2019).

49. Fawkner-Corbett, D. et al. Spatiotemporal analysis of human intestinal development at single-cell resolution. Cell 184, 810–826.e23 (2021).

50. Biton, M. et al. T Helper Cell Cytokines Modulate Intestinal Stem Cell Renewal and Differentiation. Cell 175, 1307–1320.e22 (2018).

51. Goto, N., Imada, S., Deshpande, V. & Yilmaz, Ö. H. Lymphatics constitute a novel component of the intestinal stem cell niche. bioRxiv 2022.01.28.478205 (2022) doi:10.1101/2022.01.28.478205.

52. Niec, R. E. et al. A lymphatic-stem cell interactome regulates intestinal stem cell activity. bioRxiv (2022).

53. Darling, T. K. & Lamb, T. J. Emerging Roles for Eph Receptors and Ephrin Ligands in Immunity. Front. Immunol. 10, 1473 (2019).

54. Kim, M. J. et al. PAF-Myc-Controlled Cell Stemness Is Required for Intestinal Regeneration and Tumorigenesis. Dev. Cell 44, 582–596.e4 (2018).

55. Zhang, N. et al. ID1 is a functional marker for intestinal stem and progenitor cells required for normal response to injury. Stem Cell Reports 3, 716–724 (2014).

56. Kazer, S. W. et al. Integrated single-cell analysis of multicellular immune dynamics during hyperacute HIV-1 infection. Nat. Med. 26, 511–518 (2020).

57. Strunz, M. et al. Alveolar regeneration through a Krt8+ transitional stem cell state that persists in human lung fibrosis. Nat. Commun. 11, 1–20 (2020).

58. Ravindra, N. G. et al. Single-cell longitudinal analysis of SARS-CoV-2 infection in human airway epithelium identifies target cells, alterations in gene expression, and cell state changes. PLoS Biol. 19, e3001143 (2021).

59. Gressner, O. A., Peredniene, I. & Gressner, A. M. Connective tissue growth factor reacts as an IL-6/STAT3-regulated hepatic negative acute phase protein. World J. Gastroenterol. 17, 151–163 (2011).

60. Sack, G. H., Jr. Serum amyloid A - a review. Mol. Med. 24, 46 (2018).

61. Xu, J. et al. SARS-CoV-2 induces transcriptional signatures in human lung epithelial cells that promote lung fibrosis. Respir. Res. 21, 182 (2020).

62. Doitsh, G. et al. Cell death by pyroptosis drives CD4 T-cell depletion in HIV-1 infection. Nature 505, 509–514 (2013).

63. Vignuzzi, M. & López, C. B. Defective viral genomes are key drivers of the virus–host interaction. Nature Microbiology 4, 1075–1087 (2019).

64. López, C. B. Defective viral genomes: critical danger signals of viral infections. J. Virol. 88, 8720–8723 (2014).

65. Andreatta, M. et al. Interpretation of T cell states from single-cell transcriptomics data using reference atlases. Nat. Commun. 12, 2965 (2021).

66. Svensson, V. Droplet scRNA-seq is not zero-inflated. Nat. Biotechnol. 38, 147–150 (2020).

67. Cao, Y., Kitanovski, S., Küppers, R. & Hoffmann, D. UMI or not UMI, that is the question for scRNA-seq zero-inflation. Nat. Biotechnol. 39, 158–159 (2021).

68. Ghaddar, B. & De, S. Reconstructing physical cell interaction networks from single-cell data using Neighbor-seq. Nucleic Acids Res. gkac333 (2022).

69. Giladi, A. et al. Dissecting cellular crosstalk by sequencing physically interacting cells. Nat. Biotechnol. 38, 629–637 (2020).

70. Pasqual, G. et al. Monitoring T cell-dendritic cell interactions in vivo by intercellular enzymatic labelling. Nature 553, 496–500 (2018).

71. Guidolin, D., Marcoli, M., Tortorella, C., Maura, G. & Agnati, L. F. Receptor-Receptor Interactions as a Widespread Phenomenon: Novel Targets for Drug Development? Front. Endocrinol. 10, 53 (2019).

72. Hao, Y. et al. Integrated analysis of multimodal single-cell data. Cell (2021) doi:10.1016/j.cell.2021.04.048.

73. Becht, E. et al. Dimensionality reduction for visualizing single-cell data using UMAP. Nat. Biotechnol. (2018) doi:10.1038/nbt.4314.

74. Waltman, L. & van Eck, N. J. A smart local moving algorithm for large-scale modularity-based community detection. Eur. Phys. J. B 86, 471 (2013).

75. Yip, A. M. & Horvath, S. Gene network interconnectedness and the generalized topological overlap measure. BMC Bioinformatics 8, 22 (2007).

76. Dong, J. & Horvath, S. Understanding network concepts in modules. BMC Syst. Biol. 1, 24 (2007).

77. Dimitrov, D. et al. Comparison of methods and resources for cell-cell communication inference from single-cell RNA-Seq data. Nat. Commun. 13, 1–13 (2022).

78. Wilk, A. J. et al. Charge-Altering Releasable Transporters enable phenotypic manipulation of natural killer cells for cancer immunotherapy. Blood Advances In press (2020).

79. Gierahn, T. M. et al. Seq-Well: portable, low-cost RNA sequencing of single cells at high throughput. Nat. Methods 14, 395–398 (2017).

80. Hughes, T. K. et al. Highly Efficient, Massively-Parallel Single-Cell RNA-Seq Reveals Cellular States and Molecular Features of Human Skin Pathology. bioRxiv 689273 (2019) doi:10.1101/689273.

81. Wilk, A. J. et al. A single-cell atlas of the peripheral immune response in patients with severe COVID-19. Nat. Med. 26, 1070–1076 (2020).

82. Wilk, A. J. et al. Multi-omic profiling reveals widespread dysregulation of innate immunity and hematopoiesis in COVID-19. J. Exp. Med. 218, (2021).

83. Dobin, A. et al. STAR: ultrafast universal RNA-seq aligner. Bioinformatics 29, 15–21 (2013).

84. Petukhov, V. et al. dropEst: pipeline for accurate estimation of molecular counts in droplet-based single-cell RNA-seq experiments. Genome Biol. 19, 78 (2018).

85. Butler, A., Hoffman, P., Smibert, P., Papalexi, E. & Satija, R. Integrating single-cell transcriptomic data across different conditions, technologies, and species. Nat. Biotechnol. 36, 411–420 (2018).

86. McInnes, L., Healy, J. & Melville, J. UMAP: Uniform Manifold Approximation and Projection for Dimension Reduction. arXiv [stat.ML] (2018).

87. Carter, R. A. et al. A Single-Cell Transcriptional Atlas of the Developing Murine Cerebellum. Curr. Biol. 28, 2910–2920.e2 (2018).

88. Freytag, S., Tian, L., Lönnstedt, I., Ng, M. & Bahlo, M. Comparison of clustering tools in R for medium-sized 10x Genomics single-cell RNA-sequencing data. F1000Res. 7, 1297 (2018).

89. Miller, B. C. et al. Subsets of exhausted CD8+ T cells differentially mediate tumor control and respond to checkpoint blockade. Nat. Immunol. 20, 326–336 (2019).

90. Chibueze, C. E., Yoshimitsu, M. & Arima, N. CD160 expression defines a uniquely exhausted subset of T lymphocytes in HTLV-1 infection. Biochem. Biophys. Res. Commun. 453, 379–384 (2014).

91. Agresta, L., Hoebe, K. H. N. & Janssen, E. M. The Emerging Role of CD244 Signaling in Immune Cells of the Tumor Microenvironment. Front. Immunol. 9, 2809 (2018).

92. Lopez, R., Regier, J., Cole, M. B., Jordan, M. I. & Yosef, N. Deep generative modeling for single-cell transcriptomics. Nat. Methods 15, 1053–1058 (2018).

93. Hie, B., Bryson, B. & Berger, B. Efficient integration of heterogeneous single-cell transcriptomes using Scanorama. Nat. Biotechnol. 37, 685–691 (2019).

94. Polo, J. M., Ci, W., Licht, J. D. & Melnick, A. Reversible disruption of BCL6 repression complexes by CD40 signaling in normal and malignant B cells. Blood 112, 644–651 (2008).

95. McKinlay, C. J. et al. Charge-altering releasable transporters (CARTs) for the delivery and release of mRNA in living animals. Proc. Natl. Acad. Sci. U. S. A. 114, E448–E456 (2017).

96. McKinlay, C. J., Benner, N. L., Haabeth, O. A., Waymouth, R. M. & Wender, P. A. Enhanced mRNA delivery into lymphocytes enabled by lipid-varied libraries of charge-altering releasable transporters. Proc. Natl. Acad. Sci. U. S. A. (2018) doi:10.1073/pnas.1805358115.

97. Molfetta, R., Quatrini, L., Santoni, A. & Paolini, R. Regulation of NKG2D-Dependent NK Cell Functions: The Yin and the Yang of Receptor Endocytosis. Int. J. Mol. Sci. 18, |p(2017).

98. Carlsten, M. et al. Primary human tumor cells expressing CD155 impair tumor targeting by down-regulating DNAM-1 on NK cells. J. Immunol. 183, 4921–4930 (2009).

