## Supplementary Information for "Comparative analysis of cell-cell communication at single-cell resolution"

**This PDF file includes:**

Supplementary Text
Supplementary Figures 1-14
Supplementary Tables 1-2

### SUPPLEMENTARY INFORMATION

#### Supplementary Text

<u>Selection of optimal parameters for single-cell resolution CCC analysis with Scriabin</u>

Implementing Scriabin requires users to select several parameters, particularly during ligand activity ranking as well as in the interaction program discovery workflow. Here, we discuss these parameters, their impact on obtained results, and suggestions for adjusting them based on biological and experimental considerations.

User-defined parameters for ligand activity ranking include:

1) the minimum percentage of cells expressing a ligand for that ligand to be considered expressed (% expressed ligands) – this parameter defines which ligands will be considered in Scriabin's implementation of NicheNet to identify biologically-active ligands,
2) the number of variable genes to generate the MCA embedding for calculating a cell's gene signature – this parameter reflects what genes define variation in cell state that will be captured in ligand activity ranking,
3) the distance quantile within the MCA biplot to define a cell's gene signature – this parameter defines how many genes will be used as target genes to predict ligand activities,
4) the Pearson cutoff to define an active ligand – this parameter represents the threshold of ligand activity above which the ligand will be considered biologically-active, and
5) the method for weighting the CCIM for ligand activities – this parameter determines if predicted ligand activities are considered as equally valid evidence of an interaction as expression of a cognate receptor.

To quantitatively evaluate the impact of these parameters, we leveraged data from our *in vitro* NK-B cell co-culture system as a heuristic (see **Figure 2**). In this experiment, we introduced CD40L-CD40 as an additional ligand-receptor pair edge between NK cells and B cells–thus, we evaluated which combination of parameters resulted in the most specific prediction of the CD40L-CD40 edge as differential between transfected vs. untransfected cell-cell pairs. Specifically, we applied the CCIM workflow with ligand activity ranking followed by differential expression testing using an ROC test between cells from the CD40L-CD40 transfected condition and cells from the GFP-GFP transfected condition. We define the predictive power for each ligand-receptor pair as $p = |AUC - 0.5| * 2$, and the relative predictive power for CD40L-CD40 as the difference between $p_{CD40L-CD40}$ and the average $p$ for all other ligand-receptor pairs.

We found that ligand activity ranking with every combination of parameters improved the relative log(fold-change) of predicting CD40L-CD40 as a differential ligand-receptor edge

(**Supplementary Figure 11A-C**). Using multiple regression to identify the relative impacts of each parameter on relative predictive power, we found that a lower Pearson cutoff, a higher % expressed ligands, and the use of the "sum" weighting method resulted in a higher relative prediction power. We hypothesize these trends are due to the relatively high expression of *CD40LG* in this dataset–a higher % expressed ligands prevents more lowly expressed ligands from being considered, and the "sum" weighting method allows cells with low *CD40* expression to be included in the weighting. The distance quantile and number of variable genes for cell signature calculation had little impact on the power to specifically identify CD40L-CD40 as a differential edge. Using this dataset with known ground-truth for differential CCC, we demonstrate that ligand activity ranking is relatively stable to parameter selection.

These results lead us to make the following recommendations for appropriate selection of these parameters:

1) The minimum percentage of cells expressing a ligand for that ligand to be considered expressed (default: 2.5%). We encourage users to select this parameter based on biological intuition as well as which sequencing platform was used. When ranking ligands across multiple cell types, users should set this value lower than the proportion of the rarest cell type that could be contributing to communication. Additionally, a lower value for this parameter is appropriate for droplet-based techniques relative to deeply-sequenced platforms like Smart-seq. Setting this parameter too high may result in the exclusion of ligands that are biologically important but either lowly expressed or expressed by a small subset of cells. The main issue in setting this parameter too low is an increase in computational requirements to analyze ligands that are unlikely to be biologically important. We therefore encourage users to test multiple thresholds to observe the robustness and impact of this parameter choice, and to err on the side of setting this parameter low.

2) The number of variable genes to generate the MCA embedding for calculating a cell's gene signature (default: 500). This value should be concordant with the total number of genes in a dataset that are expected to be upregulated by biologically active ligands. This parameter appears to have only a small impact on the results of the ligand activity ranking workflow.

3) The distance quantile within the MCA biplot to define a cell's gene signature (default: 0.05). This default value indicates that the top 5% of genes nearest a cell in the MCA biplot will be considered part of that cell's gene signature. Within the bounds tested (0.025 to 0.1), this parameter also appears to have only a small impact on the results on the ligand activity ranking workflow.

4) The Pearson cutoff to define an active ligand (default: 0.075). Based on benchmarking and recommendations from the author's of NicheNet[20], reasonable thresholds for defining an active ligand may lie between 0.05-0.2. We encourage users to examine the distribution of all pearson coefficients and the median pearson coefficient per cell to determine if a change from the default threshold is necessary. Generally, we have found these distributions to be right-skewed, with the right tail representing putative biological

activity. **Supplementary Figure 12** depicts an example where a Pearson threshold of 0.075 would likely capture biologically-active ligands in a system where many unknown ligands are influencing communication. In situations where a user knows *a priori* that only a subset of cells are responding to very few ligands, the pearson threshold can be raised to exclude activities the user believes may be less important.

5) The method for weighting the CCIM for ligand activities. Scriabin implements two different methods for weighting the CCIM based on predicted ligand activities. Method "product" (default) multiplies individual elements of the CCIM by scaled ligand activities that exceed the defined Pearson threshold. Method "sum" treats an active ligand prediction as equally valid evidence for an interaction as the expression of a corresponding receptor and sums receptor expression values and scaled ligand activities when calculating CCIM elements (see **Methods**). We recommend using method "product" if either of the following is true: 1) a platform with high capture/coverage (eg. Smart-seq) is used, or 2) there is a factor that decreases confidence in NicheNet's ligand-target gene linkages (for example, analysis of a non-human dataset). If neither of these considerations apply, method "sum" represents one strategy to decrease CCIM sparsity and rescue bona fide interactions that involve lowly expressed receptors.

Another important user-defined parameter in Scriabin's workflow is the soft thresholding power (softPower) used in generating the adjacency matrix for interaction program discovery. In traditional gene correlation network analyses, soft thresholding increases the degree of gene-gene connectivity required to form a module, thereby lowering the influence of spurious correlations within the similarity matrix. The authors of WGCNA have recommended utilizing the lowest softPower that results in a scale-free topology fitting index ($R^2$) of greater than 0.8[22].

Because a single ligand can interact with multiple receptors, and vice versa, the variables of a CCIM that are used to calculate the similarity matrix for WGCNA are not independent. We hypothesized that using the same soft thresholding guidelines as recommended for WGCNA may result in highly connected interaction programs where only a single ligand or receptor is represented. We thus evaluated the impact of the $R^2$ threshold on interaction program size, composition, and statistical significance.

We found that, at the standard $R^2$ = 0.8 threshold, the mean recommended softPower was 3, and this decreased as the $R^2$ threshold decreased (**Supplementary Figure 11D**). Higher $R^2$ thresholds were associated with smaller program sizes, particularly when $R^2$ > 0.5, and there was a moderate increase in the percentage of programs composed of only 1 ligand or receptor with increasing $R^2$ thresholds. Additionally, we observed that decreasing the $R^2$ threshold led to a moderate increase in the percentage of non-statistically significant programs, indicative of spurious correlations. These data indicate that an optimal $R^2$ threshold to avoid both spurious programs as well as programs composed of only a single ligand or receptor may lie between 0.5 and 0.75. The default $R^2$ for interaction program discovery is now set at 0.6.

Conceptual requirements for dataset alignment for comparative analyses of summarized interaction graphs

Comparing summarized interaction graphs from multiple samples requires that cells from different samples share a set of labels or annotations denoting what cells represent the same identity. Each identity class to be compared then requires representation from each of the samples to be compared. This annotation typically comes in the form of coarse, low-resolution labels like cluster or cell type calls. We sought to minimize the degree of agglomeration required for comparative CCC analysis by maximizing the resolution of cell type identity labels.

We hypothesized that high-resolution clustering or sub-clustering is an inadequate solution because greater transcriptional perturbation between samples necessitates lower clustering resolutions to capture representation from each sample. To illustrate this observation, we analyzed a toy dataset of peripheral blood monocytes from a longitudinal experiment (**Supplementary Figure 13**). Cells from the week 4 timepoint show a high degree of transcriptional perturbation from the other samples, visually evidenced by little overlap in low dimensional manifold embeddings. At default clustering resolution, the cluster that constitutes this sample does not contain cells from two of the samples we wish to compare. Decreasing the cluster resolution (ie. increasing the degree of agglomeration) to 0.05 improves representation of other samples but still fails to capture cells from all timepoints in each cluster. An optimal strategy for comparing summarized interaction graphs thus involves manifold alignment rather than clustering.

Scriabin's binning workflow

Identifying mutual nearest neighbors (MNNs)[25] between datasets has been implemented in several methods for dataset integration[21,36,92,93]. We reasoned that because MNNs represent pairs of cells with a shared molecular state, MNNs themselves encode single-cell resolution inter-dataset correspondences that could be generalized to high-resolution identity labels for all cells in a dataset. We refer to this process as "binning" to distinguish it from graph-based clustering. Binning with Scriabin begins by identifying MNNs between all datasets to be compared, as implemented by Seurat v3[21]. MNNs are then filtered, as we have observed that cross-cell type anchor pairs tend to have lower scores (**Supplementary Figure 14**). Cells are initially binned with the set of MNNs with which they share the highest connectivity in the shared nearest neighbor (SNN) graph, and these bin assignments are further optimized for SNN connectivity and representation of all datasets to be compared. At the end of the binning process, each cell will have a high-resolution bin identity linking it to at least one cell from all other datasets to be compared. These identities can be used as the basis for comparative analysis of CCC that maintain near single-cell resolution.

We illustrate the utility of Scriabin's binning workflow by analyzing a dataset that contains PBMCs from both a human and a mouse donor. These samples contain the same broad cell types, but there are strong species-specific phenotypic differences that prevent the same cell types from clustering together (**Supplementary Figure 14**). While we can annotate these broad cell types for the individual samples, we can only perform comparative analyses of CCC at the level of granularity that we can manually annotate. Scriabin's binning workflow enables a comprehensive alignment of the gene expression manifolds between species (**Supplementary Figure 14**).

We also evaluated the performance of Scriabin's binning strategy for comparative CCC analyses when more than two samples need to be aligned. In a toy dataset of ~14,000 cells from nine sub-datasets, Scriabin identified a total of 456 bins with a median bin size of 25 cells, maintaining near single-cell resolution (**Supplementary Figure 14**). Additionally, cells from each bin generally shared the same orthogonal reference-based cell type annotation (**Supplementary Figure 14**). For example, all plasmacytoid dendritic cells (pDCs) fell into a single bin that was composed of only pDCs. Bins whose cells did not share the same cell type annotation generally shared related annotations, for instance, intermediate and naive B cells frequently occupied the same bin (**Supplementary Figure 14**).

<u>Heterogeneity of interaction program structure and expression</u>

The discovery of interaction programs in the intestinal development dataset[49] revealed several programs that were significantly up- or down-regulated between anatomical sampling locations, reflecting either changes in the magnitude of program expression or a shift in the proportions of cell types expressing the program (**Supplementary Figure 8**). We also hypothesized that interaction program structure may vary between samples even if the expression magnitude of the program does not change. Thus, we calculated intramodular connectivity scores for each gene in each sampling location analyzed. We identified ligand-receptor pairs in non-differentially expressed interaction programs that had differential intramodular connectivity between sampling locations (**Supplementary Figure 8**). For example, the *EFNB2* - *EPHB4* ligand-receptor pair has higher intramodular connectivity in hindgut than other anatomical locations, and the hindgut is the location that expresses the highest level of both of these genes (**Supplementary Figure 8**). This indicates that, in the hindgut, when *EFNB2* and *EPHB4* are expressed, they are co-expressed along with other ligand receptor pairs in this interaction program. Collectively, this analysis provides an example of the heterogeneity, specificity, and nuanced co-expression patterns that can be revealed through the scalable discovery of interaction programs in atlas-scale datasets.

<u>Validation of Scriabin through transfection of exogenous CCC edges</u>

To provide additional evidence that Scriabin recovers differentially-expressed and biologically active CCC edges, we designed an experiment where we would force overexpression of a specific ligand-receptor pair, co-culture the cells, and then profile the co-cultured cells by scRNA-seq. Thus, we transfected isolated human NK cells with CD40L-encoding mRNA and isolated human B cells with CD40-encoding mRNA, followed by 12 hour co-culture and single-cell transcriptional profiling by Seq-Well[79]. We chose to profile interactions between NK cells and B cells because at baseline these cell types do not express a large number of cognate ligand-receptor pairs. We selected the CD40L-CD40 interaction because the transcriptional response downstream of CD40 in B cells is well-characterized. Further, we decided to co-culture the cells for 12 hours because this is the time at which the transcriptional response to CD40 signaling peaks[94]. Finally, we transfected the isolated NK and B cells with mRNA complexed with Charge-Altering Releasable Transporters, a class of cationic di-block oligomers that degrade into neutral small molecule metabolites after delivering anionic cargo to a cell, both facilitating cargo release and avoiding toxicity associated with persistent cations[95,96]. We chose CARTs because we have recently shown that, unlike lipofection and electroporation, CARTs transfect resting primary immune cells with high efficacy while causing minimal reconfiguration of cell surface proteome[78], making them suitable delivery vehicles for specific manipulation of cell phenotype with few off-target effects. We also elected to transfect cells with a CART initiated with a difluoroboron-β-diketonate (BDK) fluorophore so that transfected cells could be easily distinguished by flow cytometry.

Flow cytometric analysis at the start of co-culture indicated successful transfection, with 21% of B cells expressing CD40 (with a higher mean fluorescent intensity and compared to 8% at baseline), and 13.5% of NK cells expressing CD40L (compared to 1% at baseline; **Supplementary Figure 4**). After co-culture, B cells had upregulated CD40, consistent with known outcomes of CD40L/CD40 signaling[20], and NK cells had lost expression of CD40L, suggesting internalization of CD40L post-ligation, a well-established process for ligands and receptors expressed by NK cells (**Supplementary Figure 4**)[97,98]. scRNA-seq profiling by Seq-Well indicated that cells expressed the exogenous mRNAs expected from the individual co-cultures (**Supplementary Figure 4**).

### Supplementary Figures

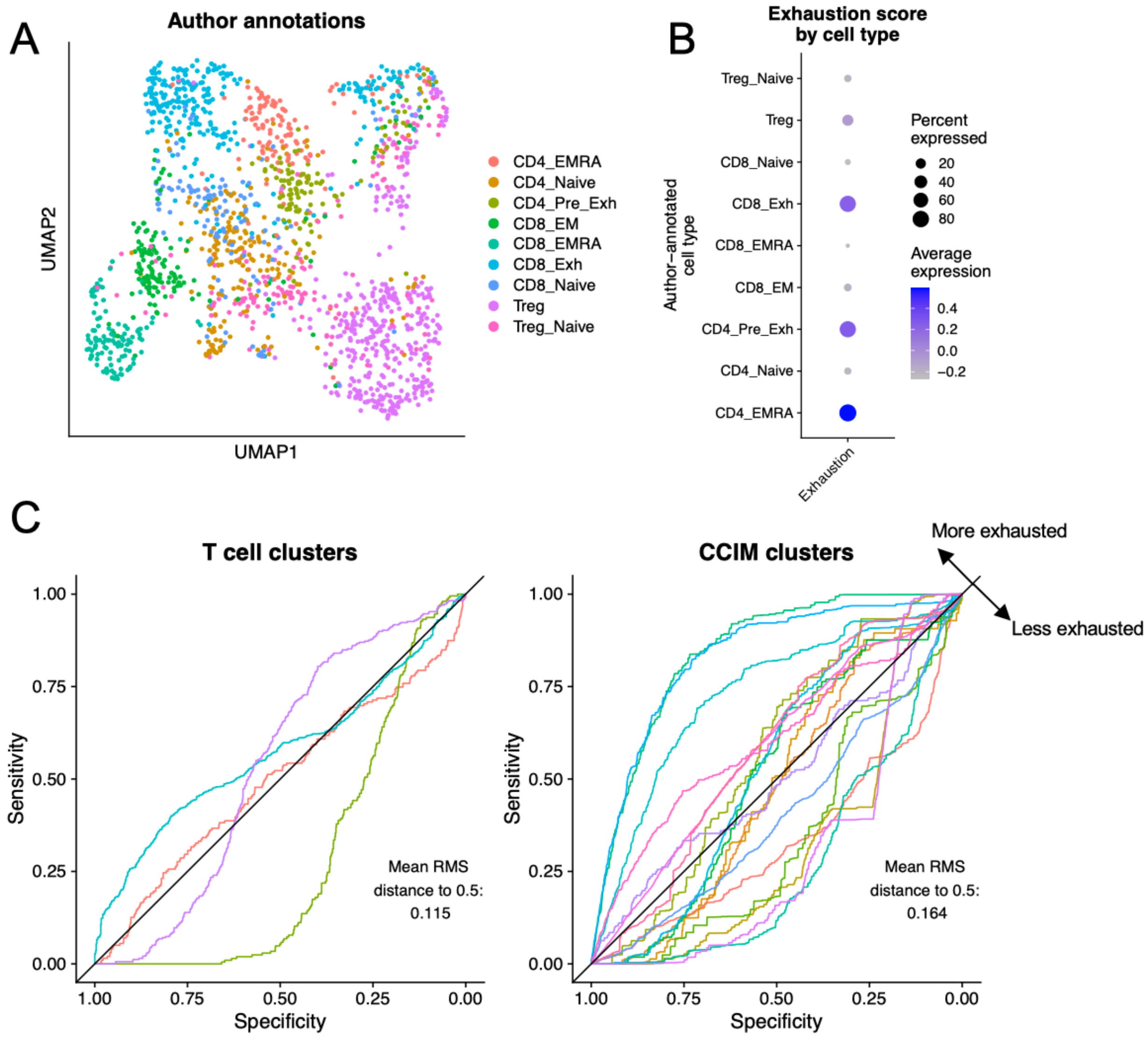


**Supplementary Figure 1: Additional analyses of exhausted intratumoral SCC T cells. A)** UMAP projection of all T cells from the dataset published by Ji, et al.[29], colored by author-annotated T cell subtype. **B)** Dot plot depicting average and percent expression of the exhaustion signature score by author-annotated T cell subtypes. **C)** ROC curves depicting the ability of each cluster from the single-cell T cell object (left) or Scriabin generated T cell-*CD1C*$^{+}$ DC CCIM (right) to be classified as exhausted or non-exhausted. Each line corresponds to a single cluster. The diagonal black line corresponds to an AUC = 0.5, where there is no predictive power of classification. AUC = 0, the cluster can be perfectly classified as non-exhausted; AUC = 1, the cluster can be perfectly classified as exhausted.

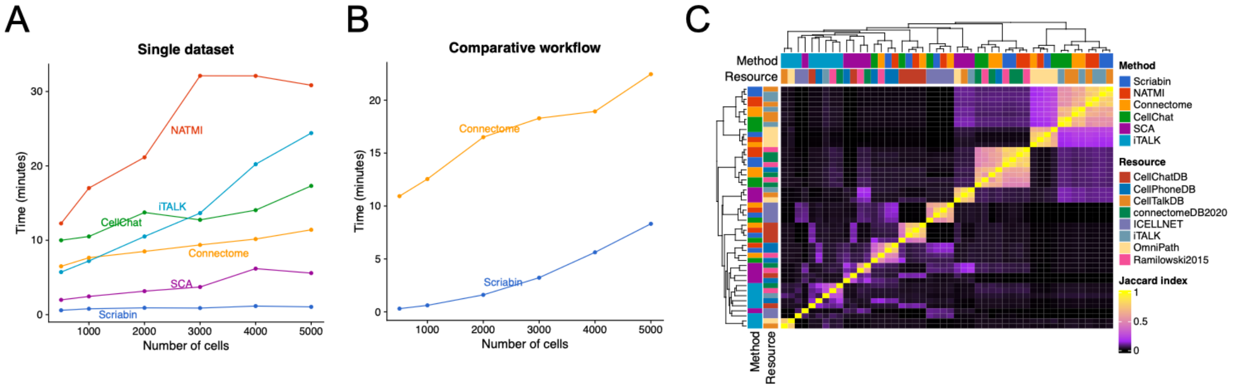


**Supplementary Figure 2: Comparison of Scriabin to agglomerative CCC analysis techniques. A)** Runtime of Scriabin and five published CCC methods on the 10X PBMC 5k dataset. For each dataset size, the dataset was randomly subsampled to the indicated size and the same subsampled dataset was used for all methods. **B)** Runtime of Scriabin and Connectome comparative workflows. The 10X PBMC 5k and 10k datasets were merged into a single dataset which was subsampled as in (**A**), and the comparative workflows performed between cells from the 5k vs. 10k dataset. **C)** Jaccard index heatmap depicting the degree of overlap in the top 1,000 ligand-receptor CCC edges from each method-resource pair.

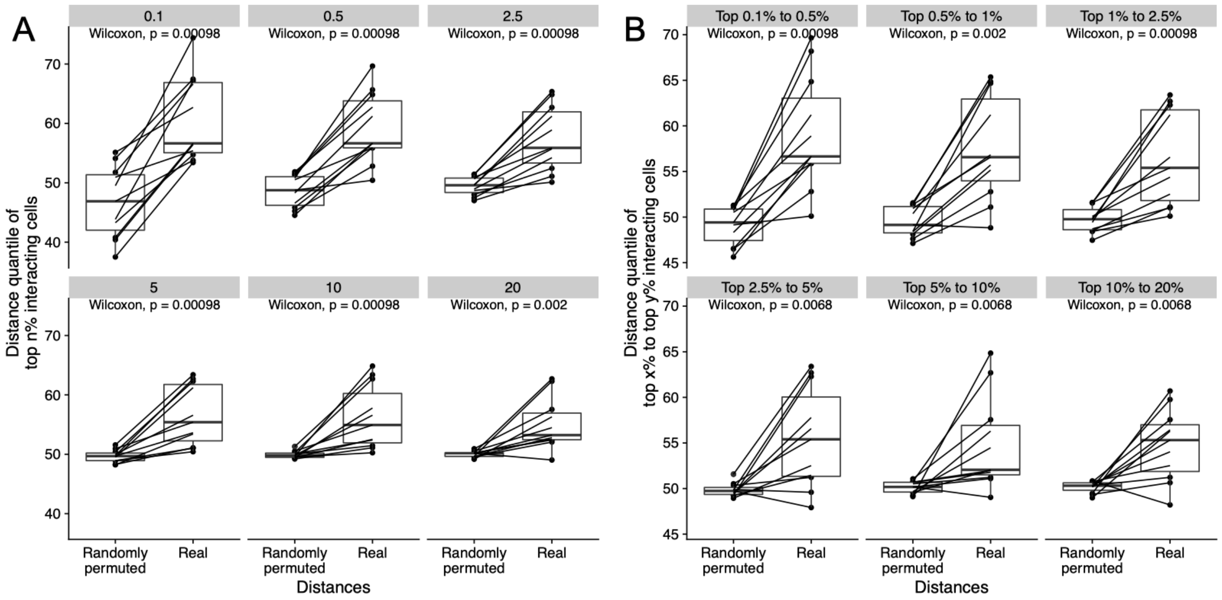


**Supplementary Figure 3: Scriabin's predictions of spatial features are robust at multiple distances cutoffs.** The procedure described in **Figure 2F** was repeated for 11 datasets, and the median distance quantile of a percentile of the most highly interacting cell-cell pairs was calculated using real cell distances relative to randomly permuted cell distances. **A)** Each facet shows the median distance quantile of the top 0.1%, 0.5%, 2.5%, 5%, 10%, and 20% most highly interacting cell-cell pairs. **B)** Each facet shows the median distance quantile of cell-cell pairs within the range of interaction quantile shown. In each facet, an exact two-sided p-value from the Wilcoxon rank-sum test is shown.

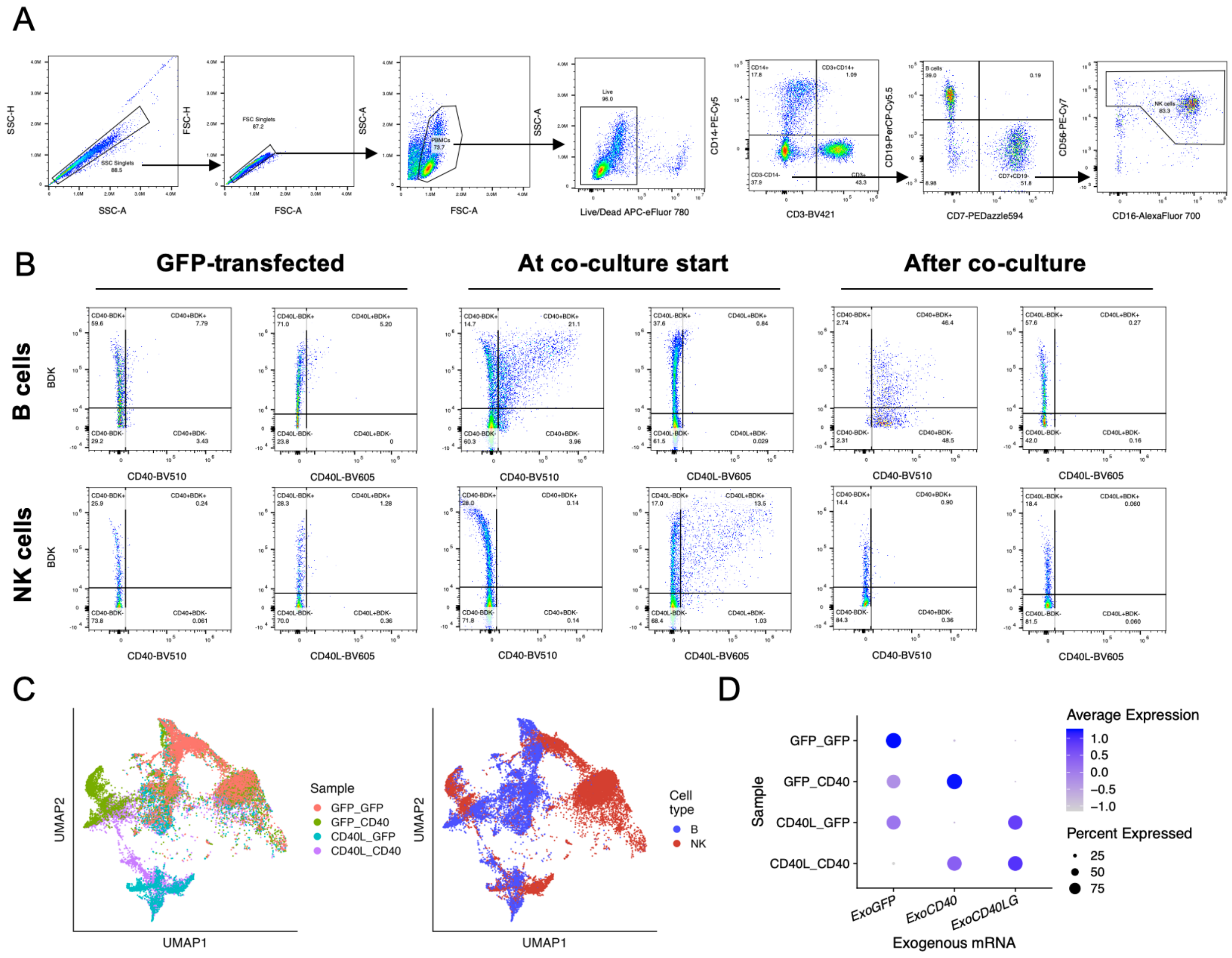


**Supplementary Figure 4**: **Flow cytometry and transcriptional analysis of B and NK cell transfection and co-culture. A)** Flow cytometry gating scheme used to identify B and NK cells. **B)** Scatter plots of flow cytometry data showing expression of CD40 and CD40L by B cells and NK cells. Left: B cells and NK cells transfected with GFP-encoding mRNA at start of co-culture. Middle: B cells transfected with CD40-encoding mRNA and NK cells transfected with CD40L-encoding mRNA at start of co-culture. Right: B cells transfected with CD40-encoding mRNA and NK cells transfected with CD40L-encoding mRNA at end of co-culture. For (**A**-**B**), percentages of the parent gate are shown for each gate. **C)** UMAP projections of full dataset colored by cell condition of origin (left) or annotated cell type (right). **D)** Dot plot depicting average and percent expression of exogenous mRNAs in the four co-cultures.

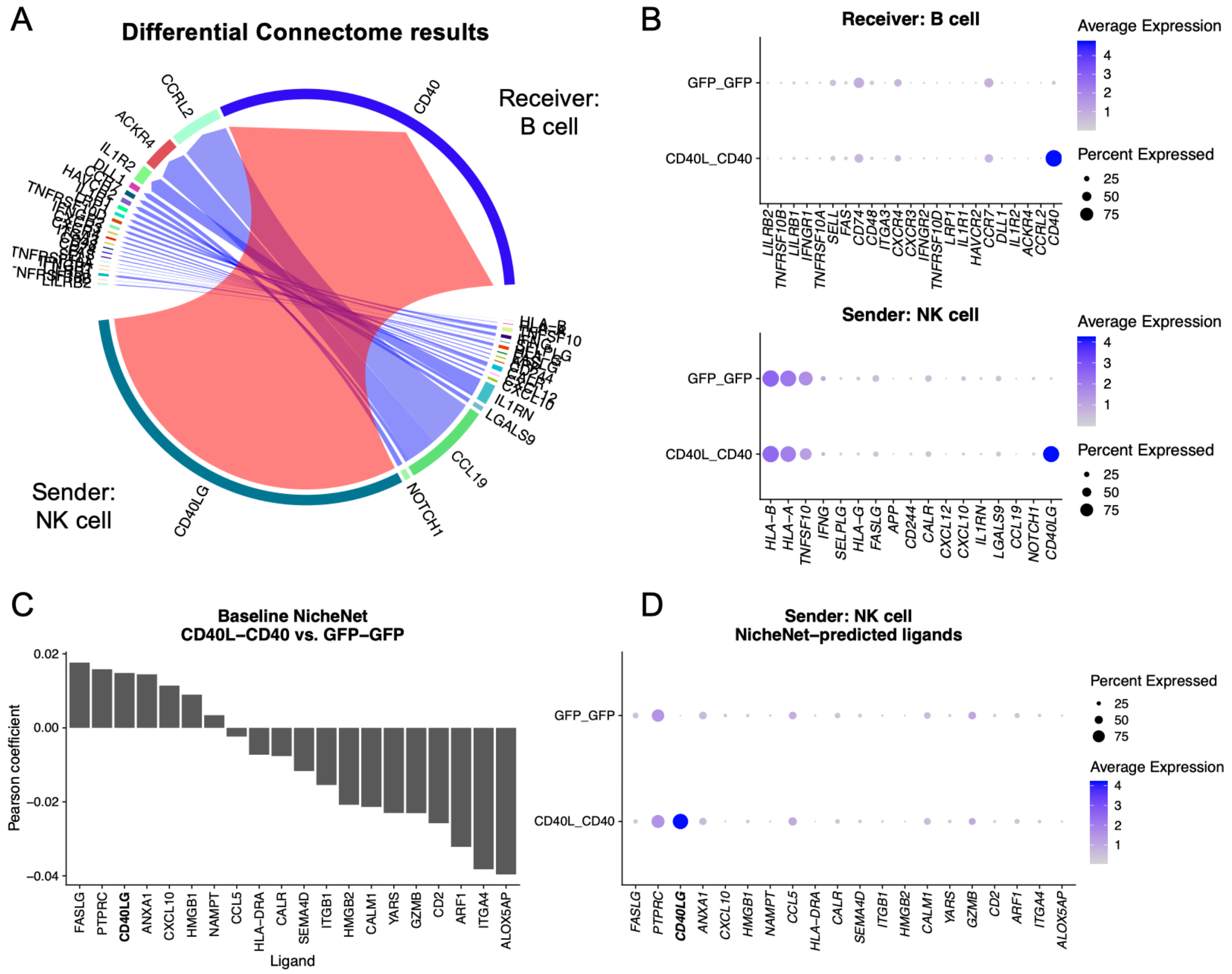


**Supplementary Figure 5: Analysis of CCC between *CD40LG*-transfected NK cells and *CD40*-transfected B cells with Connectome and NicheNet. A)** Circos plot summarizing Connectome's[38] results of significantly differentially-expressed ligand-receptor pair edges between the CD40LG-CD40 transfected condition (shades of red) and GFP-GFP transfected condition (shades of blue). CCC is analyzed between ligands expressed by sender NK cells (bottom) and receptors expressed by receiver B cells (top). **B)** Dot plot depicting percentage and average expression of differentially-expressed receptors by B cells (top) and ligands by NK cells (bottom) returned by Connectome's DifferentialConnectome workflow. **C)** NicheNet[20] was applied to predict ligand activities in B cells between the CD40LG-CD40 transfected condition and the GFP-GFP transfected condition. The bar plot depicts pearson coefficient outputs of NicheNet for this analysis. **D)** Dot plot depicting percentage and average expression of potentially-active ligands shown in (**C**) by NK cells.

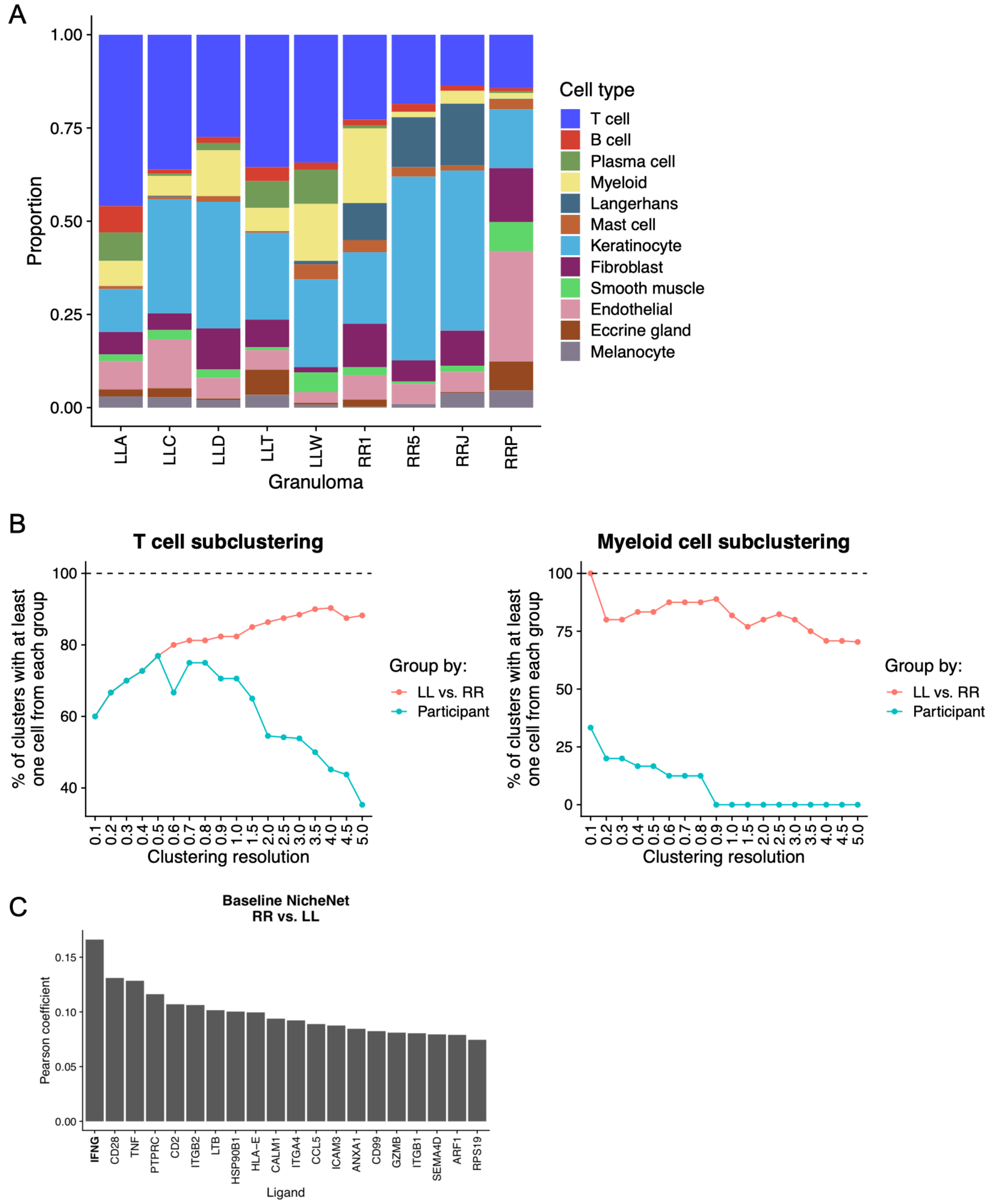


**Supplementary Figure 6**: **Additional analyses of the scRNA-seq dataset of leprosy granulomas. A)** Bar graph depicting cell proportions per granuloma in the dataset of Ma, et al[41]. Author-provided cell type annotations are used for analysis. **B)** Subclustering resolutions of T cells (left) and myeloid cells (right) required for comparative CCC analysis by agglomerative methods. Pink bars indicate the percentage of subclusters containing at least one cell from an LL granuloma and one cell from an RR granuloma. Blue bars indicate the percentage of subclusters containing at least one cell from all nine analyzed granulomas. **C)** NicheNet[20] was applied to predict ligand activities in myeloid cells between RR granulomas relative to LL granulomas. The bar plot depicts pearson coefficient outputs of NicheNet for this analysis.

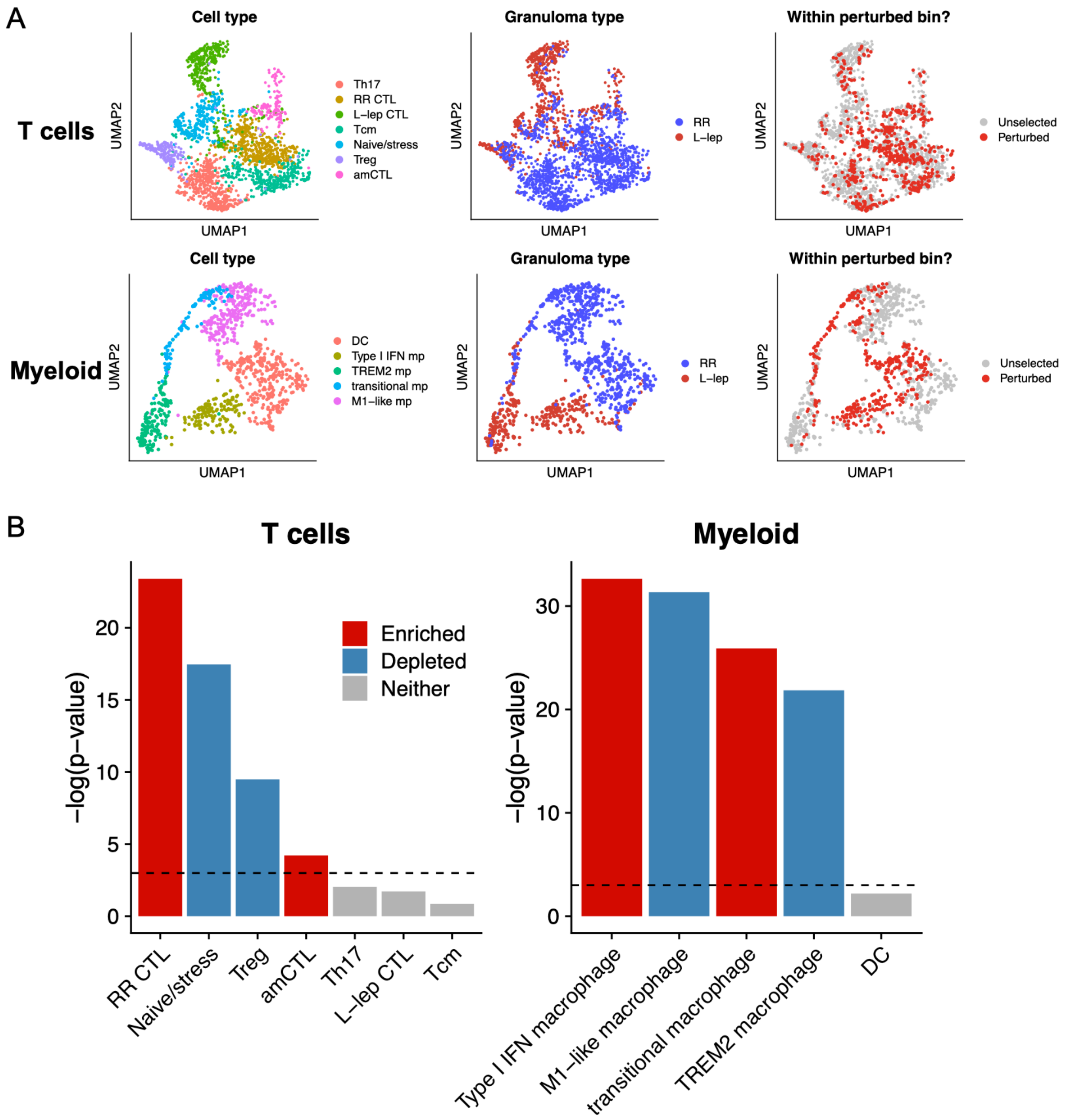


**Supplementary Figure 7: Enrichment of T and myeloid cells with perturbed CCC in RR vs. LL granulomas in T and myeloid cell subclusters. A)** UMAP projections of T cells (top) and myeloid cells (bottom) colored by author-generated subcluster cell type annotation (left), granuloma type (middle), or if the cell falls into a cluster 2 perturbed bin (right; see **Figure 3F**). **B)** We applied a binomial test to determine if cells from a cluster 2 perturbed bin were significantly enriched or depleted in any T cell or myeloid cell subcluster. The bar plot depicts the -log(p-value) of the exact binomial test. When $p < 0.05$, the bars are colored to indicate if perturbed cells are either enriched (red) or depleted (blue) from the cluster. The dotted line indicates the point at which $p = 0.05$.

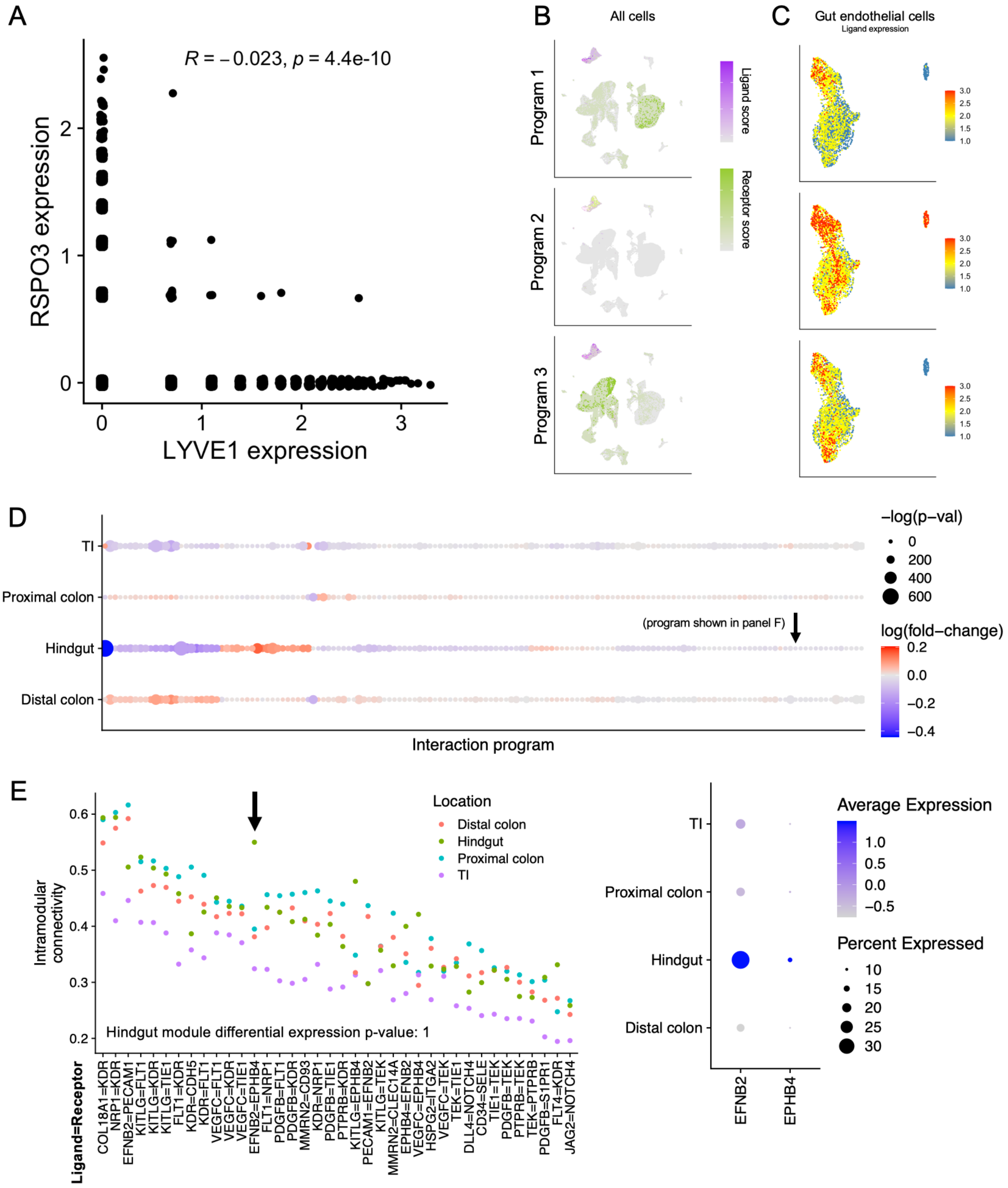


**Supplementary Figure 8: Co-expressed interaction programs in intestinal development.**
**A**) Scatter plot depicting expression of LEC marker *LYVE1* and *RSPO3*. **B)** UMAP projections of ligand (shades of purple) or receptor (shades of green) expression in 3 gut endothelial cell-specific modules. **C)** UMAP projection of gut endothelial cells colored by expression of ligands in the interaction programs depicted in (**B**). **D)** Dot plot depicting the expression

fold-change and Bonferroni-corrected Wilcoxon rank-sum test 2-sided p-values of interaction program expression in each anatomical location. **E)** Intramodular connectivity scores for each ligand-receptor pair in each anatomical location for the module indicated by the arrow in (**D**). The black arrow in (**E**) indicates the genes whose average and percent expression are plotted to the right.

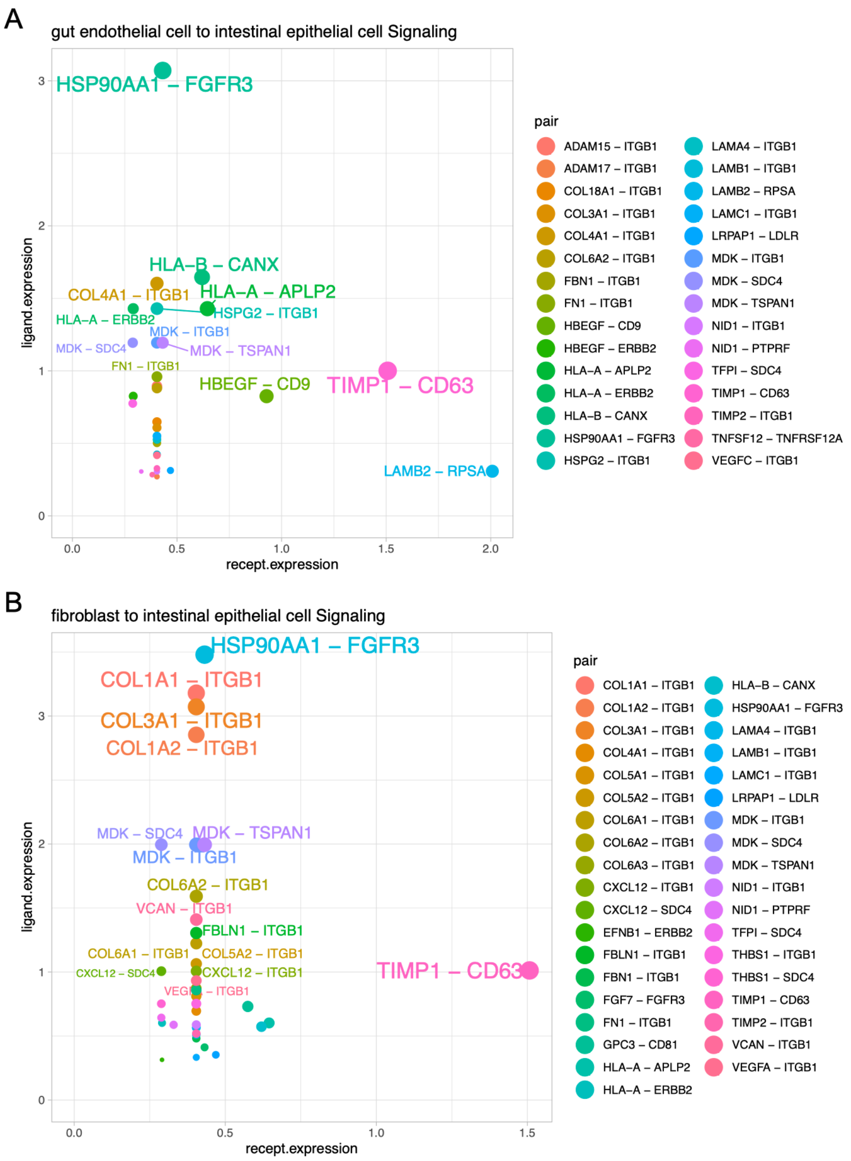


**Supplementary Figure 9. Agglomerative analysis of CCC in human intestinal development dataset.** Connectome[38] was used to analyze CCC in the human intestinal development dataset[49] using author-annotated cell types for aggregation. Results are plotted for communication between gut endothelial cells (senders) and intestinal epithelial cells (receivers; **A**) or between fibroblasts (senders) and intestinal epithelial cells (receivers; **B**).

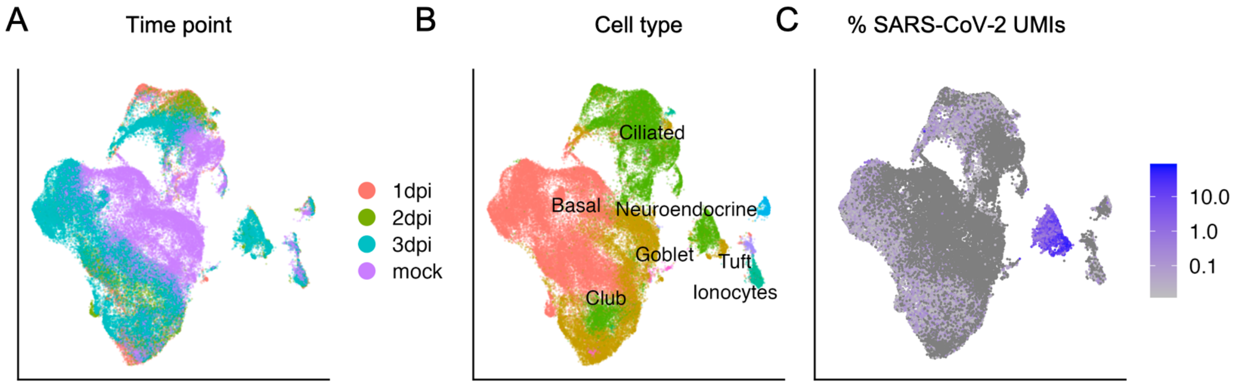


**Supplementary Figure 10. scRNA-seq dataset of SARS-CoV-2 infected HBECs.** UMAP projections of 64,008 cells from the dataset published by Ravindra, et al.[58] colored by time point (**A**), annotated cell type (**B**), or the percentage of UMIs per cell of SARS-CoV-2 origin (**C**).

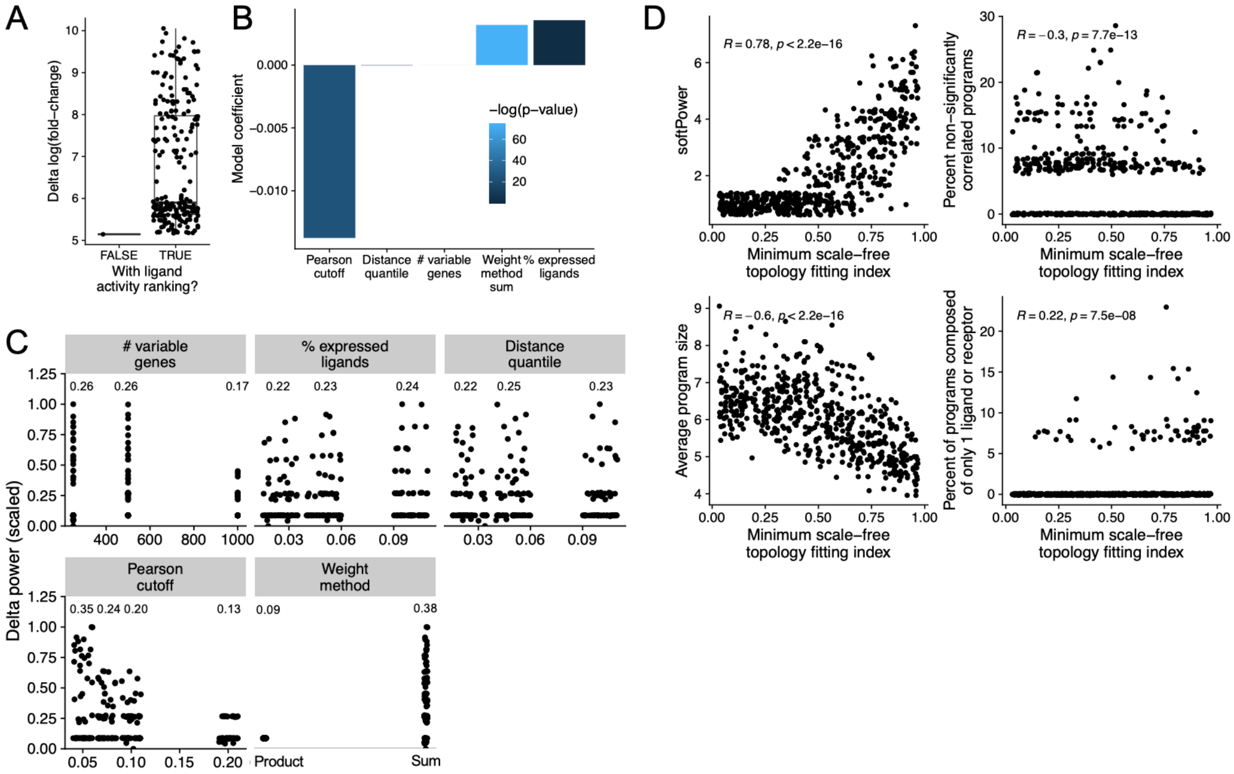


**Supplementary Figure 11: Parameter tuning for ligand activity ranking and interaction program discovery workflows. A-C)** 217 different parameter combinations were used to analyze CCC between NK cells transfected with CD40L-encoding mRNA and B cells transfected with CD40-encoding mRNA. Ligand activity-weighted CCIMs were calculated from each of these combinations and differential expression testing performed to identify which parameter combinations returned CD40L-CD40 as a differential edge with the highest specificity. **A)** Box plot depicting the difference between the log(fold-change) for CD40L-CD40 and the mean log(fold-change) for all other ligand-receptor pairs, with and without application of ligand activity ranking. **B)** Scatter plots depicting relative predictive power for the CD40L-CD40 edge for all combinations of ligand ranking parameters. The mean for each parameter is shown within the plot. **C)** β coefficients and p-values from multiple regression analysis modeling the impact of each ligand ranking parameter on relative predictive power for the CD40L-CD40 edge. **D)** The interaction program discovery workflow was repeated on 35 random subsamples of the inDrop panc8 dataset[21,34], using 19 different $R^2$ thresholds to define the appropriate softPower parameter. Scatter plots depict association between $R^2$ threshold and (clockwise from top left): recommended softPower, percentage of identified programs that failed significance testing, percentage of programs composed of only 1 ligand or receptor, and the average number of ligands and receptors composing a program.

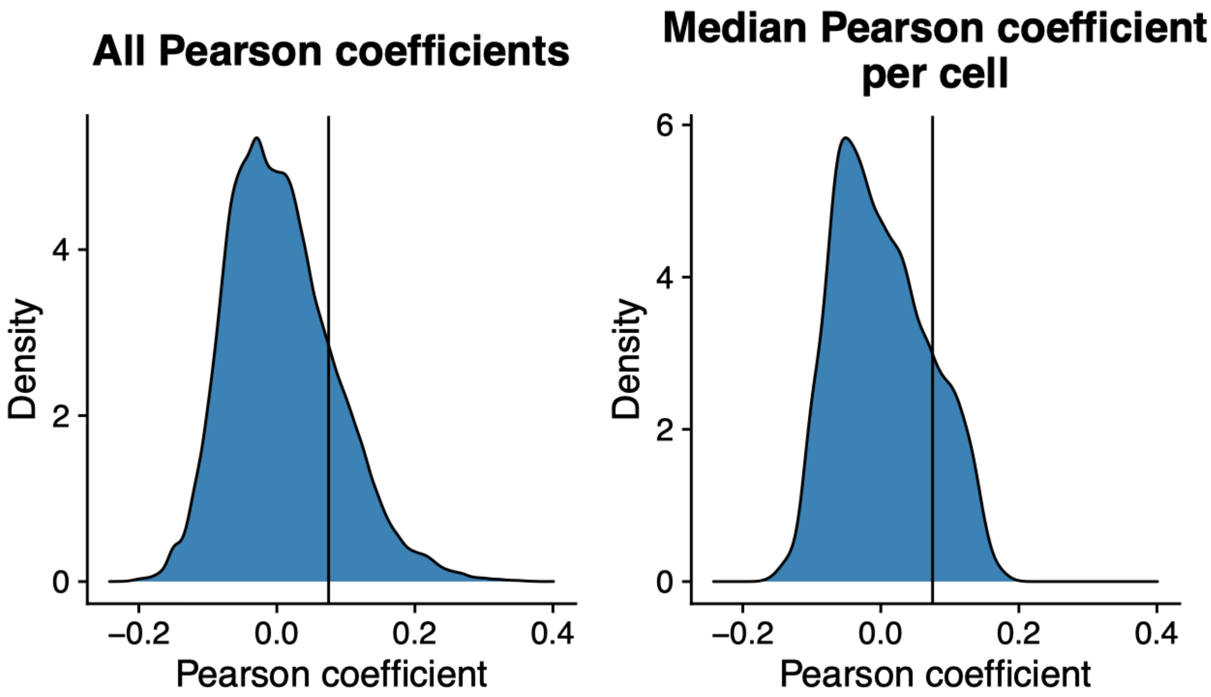


**Supplementary Figure 12. Example ligand activity distributions to aid in selection of the appropriate Pearson coefficient threshold.** Generally, ligand activity coefficients form a right-skewed distribution, similar to the distributions shown here. The right tails of these distributions represent the putative biological activity and are the coefficients that should be used for CCIM weighting. We therefore encourage users to consider the number of ligands that are expected to display biological activity and the number of cells that are expected to have downstream signaling induced by those ligands. If there are very few ligands expected to be biologically active, and only a subset of cells responding to them, this threshold should be increased to include less of the right tail of the distribution.

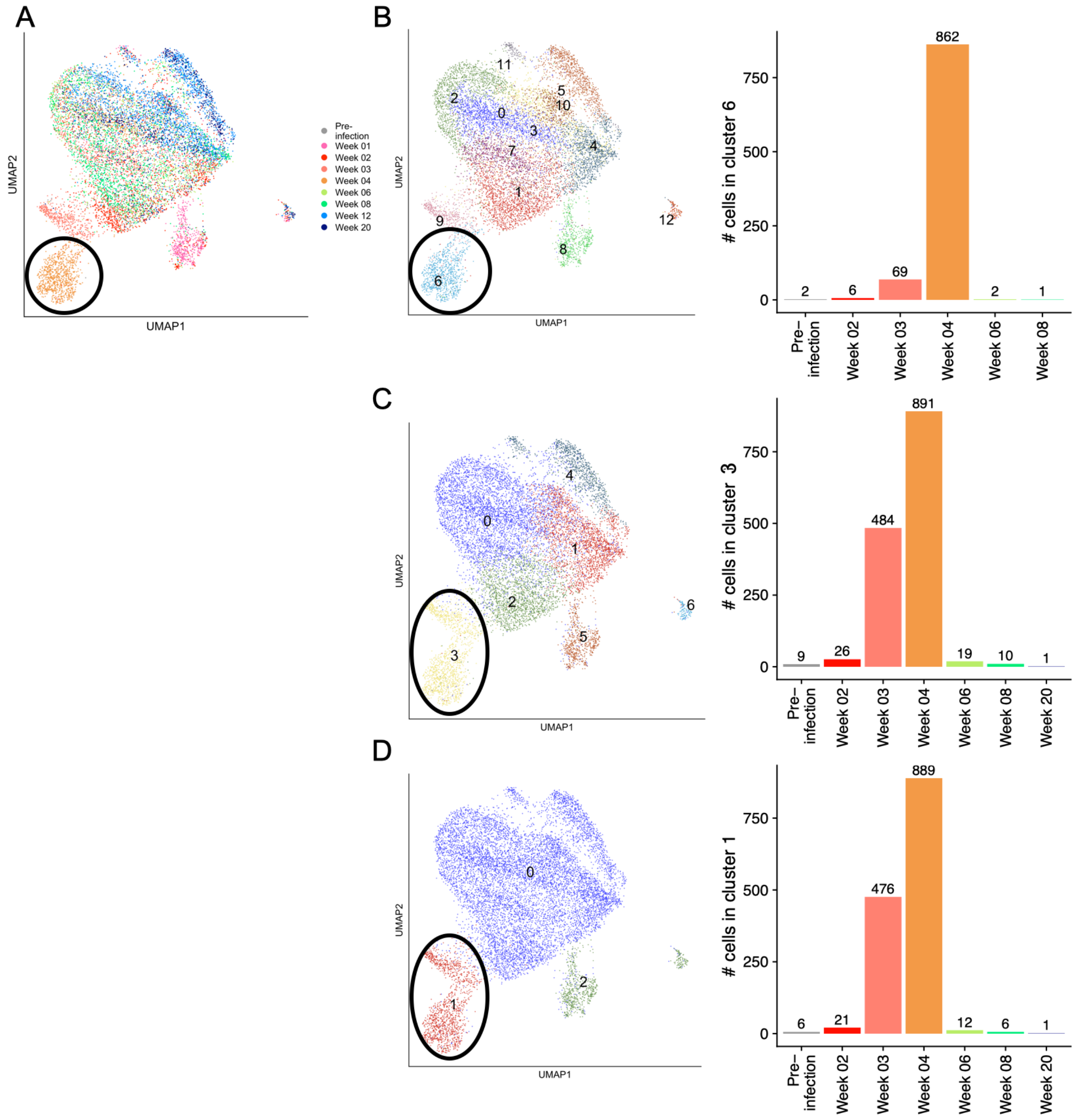


**Supplementary Figure 13: Highly perturbed samples require a higher degree of aggregation for dataset alignment**. A toy dataset of peripheral blood monocytes from a longitudinal dataset was analyzed. **A)** UMAP projection colored by time point. **B-D)** UMAP projections (left) colored by cluster identity, and bar plot depicting per timepoint cluster membership in the cluster principally occupied by sample Week 04 (right). Cluster resolutions: 1 (default, **B**), 0.3 (**C**), 0.05 (**D**).

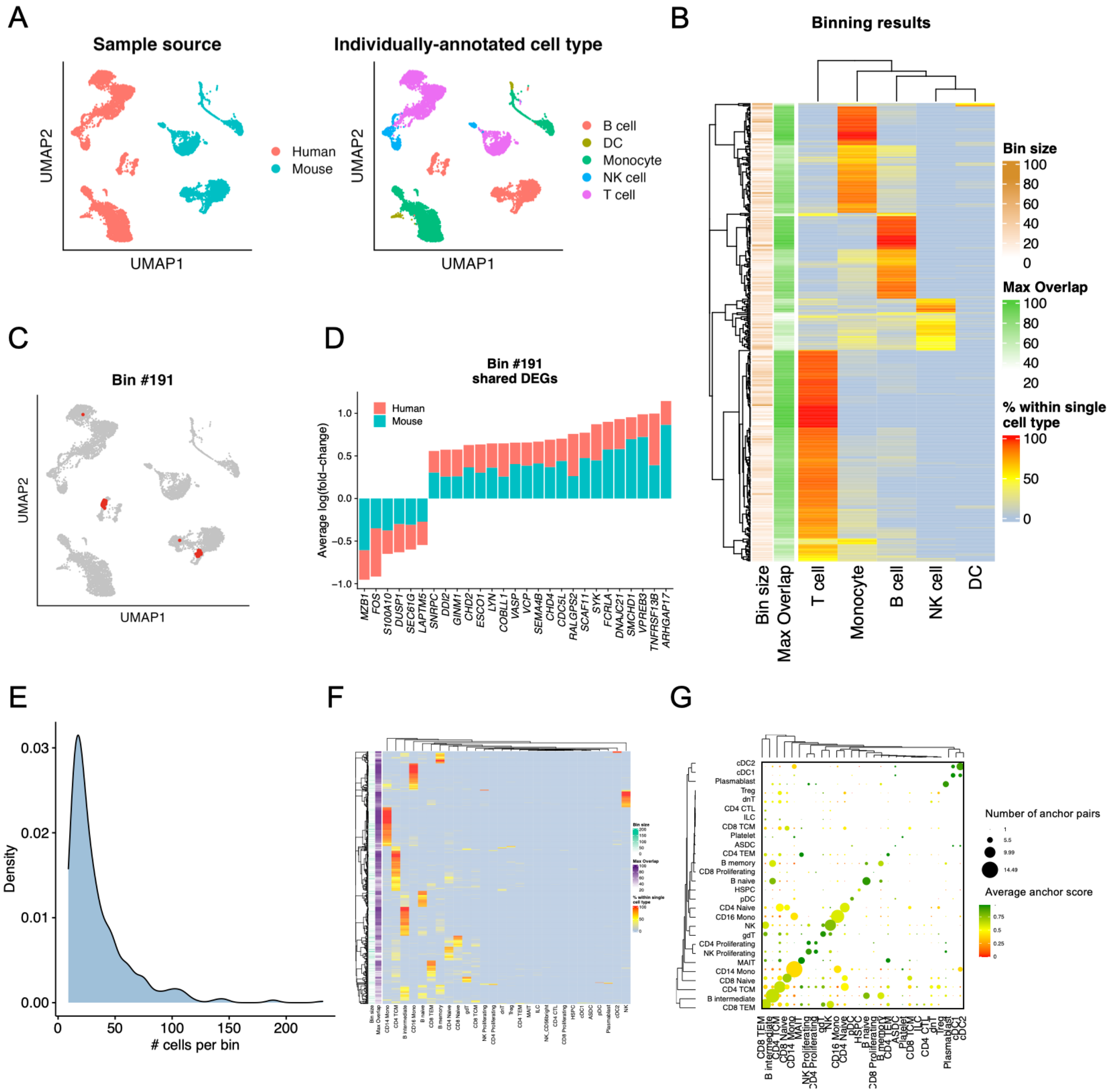


**Supplementary Figure 14: Robustness analysis of Scriabin's binning workflow. A-D)** Mouse and human PBMC scRNA-seq datasets from 10X Genomics were analyzed. **A)** UMAP projections of mouse and human PBMCs colored by the sample of origin (left) and by manually-annotated cell types (right). **B)** Heatmap depicting overlap between bin identity and cell type annotations. Each row sums to 100%, and the annotations at left show the number of cells within each bin and maximum degree of overlap of each bin with a given cell type identity (ie. the highest value in each row). **C)** UMAP projection highlighting cells in bin #191. **D)** Bar plot depicting differentially-expressed genes in bin #191 relative to other B cells shared between the human and mouse cells in bin #191. Differential expression tests were run individually for human and mouse cells. **E-G)** A toy dataset of ~14,000 peripheral blood mononuclear cells (PBMCs) from nine sub-datasets was analyzed. **E)** Density plot depicting the number of cells in each bin. The median bin size in this analysis is 25 cells. **F)** As in (**B**) An SNN graph was used to assess cell-cell connectivity for the binning workflow. Cell type annotations are transferred from a reference dataset and are thus orthogonal to the data used to generate the bins. **G)** Dot plot depicting the cell type annotations and scores for the anchor pairs used to generate the bins depicted in (**F**).

**Supplementary Tables**

| **Sample** | **T cells** | **Myeloid cells** | **Total** |
|---|---|---|---|
| LLA | 1377 | 201 | 2996 |
| LLC | 380 | 56 | 1052 |
| LLD | 732 | 328 | 2671 |
| LLT | 320 | 56 | 902 |
| LLW | 337 | 150 | 983 |
| RR1 | 1153 | 1014 | 5062 |
| RR5 | 690 | 52 | 3721 |
| RRJ | 213 | 54 | 1574 |
| RRP | 317 | 35 | 2216 |

**Supplementary Table 1**. Sample sizes for analysis of leprosy granuloma dataset published by Ma, et al.

| **Marker** | **Fluorophore** | **Clone** | **Source** |
|---|---|---|---|
| CD3 | BV421 | OKT3 | BioLegend |
| CD7 | PE-Dazzle594 | CD7-6B7 | BioLegend |
| CD14 | PE-Cy5 | 61D3 | ThermoFisher |
| CD16 | AlexaFluor700 | 3G8 | BioLegend |
| CD19 | PerCP-Cy5.5 | HIB19 | BD Biosciences |
| CD56 | PE-Cy7 | HDC56 | BioLegend |
| CD40 | BV510 | 5C3 | BioLegend |
| CD40L | BV605 | 24-31 | BioLegend |

**Supplementary Table 2**. Antibodies used for flow cytometric analysis.
